# The Role of the Ventrolateral Anterior Temporal Lobes in Social Cognition

**DOI:** 10.1101/2021.09.10.459496

**Authors:** Eva Balgova, Veronica Diveica, Jon Walbrin, Richard J. Binney

## Abstract

A key challenge for neurobiological models of social cognition is to elucidate whether brain regions are specialised for that domain. In recent years, discussion surrounding the role of anterior temporal regions epitomises such debates; some argue the anterior temporal lobe (ATL) is part of a domain-specific network for social processing, while others claim it comprises a domain-general hub for semantic representation. In the present study, we used ATL-optimised fMRI to map the contribution of different ATL structures to a variety of paradigms frequently used to probe a crucial social ability, namely ‘theory of mind’ (ToM). Using multiple tasks enables a clearer attribution of activation to ToM as opposed to idiosyncratic features of stimuli. Further, we directly explored whether these same structures are also activated by a non-social task probing semantic representations. We revealed that common to all of the tasks was activation of a key ventrolateral ATL region that is often invisible to standard fMRI. This constitutes novel evidence in support of the view that the ventrolateral ATL contributes to social cognition via a domain-general role in semantic processing, and against claims of a specialised social function.

## Introduction

The anterior temporal lobe (ATL) plays a crucial role in support of social cognition (Frith and Frith 2003, 2010; Olson et al. 2013; Binney and Ramsey 2020). Damage to the ATL results in profound and wide-ranging socio-affective deficits in both primates and humans (Klüver and Bucy 1937; Terzian and Dalle Ore 1955; Edwards-Lee et al. 1997; Kumfor and Piguet 2012; Kumfor et al. 2013, 2017; Irish et al. 2014; Binney et al. 2016). Amongst neurotypical samples, the findings of functional neuroimaging studies suggest an almost ubiquitous involvement in the high-level processing of faces and emotions (Wong and Gallate 2012; Avidan et al. 2014; Collins and Olson 2014; Collins et al. 2016; Ramot et al. 2019), as well as in more abstracted forms of social processing, such as moral cognition and mental state attribution (also known as theory of mind) (Moll et al. 2005; Schurz et al. 2014; Molenberghs et al. 2016; Diveica et al. 2021).

Despite this, across various neurocognitive frameworks of the ‘social brain’, there is no firm consensus regarding the nature of the function that the ATL performs (for a comprehensive review, see Olson et al. 2007; 2013). There are likely two main drivers for this. First, at a glance, it might be difficult to identify a common cognitive process that connects the various social and emotional tasks that implicate the ATL (Olson et al. 2013; Binney and Ramsey 2020). Second, inconsistent definitions of what anatomy it is that constitutes the ATL have greatly contributed to a lack of clarity regarding the locations at which overlap, and divergence of function seemingly occurs. From one perspective, the term “ATL” refers to all cortex comprising the anterior half of the temporal lobe (Binney et al. 2010; Rice, et al. 2015; Binney et al. 2016; Rice et al. 2018), and therefore a large area potentially comprising a number of functionally distinct subregions. However, it has at times been used to more specifically refer to the temporal polar cortex, and the limited boundaries of Brodmann’s area 38 (e.g., Ross and Olson 2010; Simmons et al. 2010). Therefore, the primary aims of the present study were to contribute to a more complete description of the ATL subregions engaged in service of social cognitive tasks, and to advance understanding of the nature of their function.

One account of the ATL’s role in social tasks is that it stores mental scripts, or schema, that are formed out of prior experiences, and provide a wider context for understanding social interactions (Frith and Frith 2003; Gallagher and Frith 2003). However, until more recently, there has been a lack of direct evidence to support this hypothesis. Moreover, it is unclear as to what extent the proposed social functions played by ATL subregions are distinct from that of more general declarative memory systems. Recent proposals have specifically associated the role of some ATL subregions with the retrieval of social conceptual knowledge, which is posited as a subtype of semantic memory (Zahn et al. 2007; Olson et al. 2013; Binney and Ramsey 2020). Semantic memory is a term used to refer to a long-term store of general conceptual-level knowledge that is involved in transforming sensory inputs into meaningful experiences, and it underpins the ability to recognize and make inferences about objects, people, and events in our environment (Lambon Ralph et al. 2017). Social conceptual knowledge has been defined more distinctively as person-specific knowledge (Simmons et al. 2010), but also knowledge about interpersonal relationships, social behaviours, and of more abstract social concepts such as *truth* and *liberty* (Zahn et al. 2007; Olson et al. 2013). The claim that the ATL is engaged in retrieving this type of information during social tasks is supported by a functional neuroimaging study that reveals ATL activation that is common to both a social attribution task and a task involving semantic relatedness judgments about socially relevant concepts (Ross and Olson 2010).

Moreover, it has been suggested that social conceptual knowledge could have a special, or even privileged status over other categories of semantic information (Zahn et al. 2007; Olson et al. 2013). Indeed, one influential account of the ATL, the *social knowledge hypothesis*, states that the dorsolateral portion (including the anterior superior and middle temporal gyri) is selectively involved in processing social concepts (Olson et al. 2013) This account has some overlap with the *social information processing account* which claims that the dorsolateral ATL activates to social concepts as part of a larger domain-specific network involved in social information processing (Simmons et al. 2010; Persichetti et al. 2021). Indeed, some fMRI studies have demonstrated a greater response of dorsolateral ATL subregions when semantic judgments made on socially relevant stimuli are compared to similar judgments made on non-social stimuli (Zahn et al. 2007; Ross and Olson 2010; Binney et al. 2016; Rice et al. 2018). Further, proponents of the social knowledge hypothesis argue against a more general role of ATL subregions in semantic processing, and point to the fact that a variety of socially-relevant tasks and stimuli reliably activate the ATL, whereas the majority of functional imaging studies of general semantic processing do not (Olson et al. 2007, 2013; Simmons and Martin 2009; Simmons et al. 2010).

However, the ATL is strongly implicated in general semantic processing on the basis of decades of neuropsychological data (Patterson et al. 2007) and a growing body of brain stimulation and electrophysiological studies, as well as functional neuroimaging studies that take special measures to address signal dropout and distortion within this region (Binney et al. 2010; Visser et al. 2010; Visser et al. 2010; Lambon Ralph et al. 2017). A critical issue, therefore, is how it is possible to reconcile these two sets of observations within a single unified theory of ATL function.

When broadly defined as the anterior half of the temporal lobe, the ATL is comprised of a substantial volume of cortex, amongst which there are numerous subdivisions identifiable on the basis of morphology, cytoarchitecture and connectivity (Ding et al. 2009; Binney et al. 2012; Pascual et al. 2015), and it is highly plausible that within it there are either distinct functional parcels (Perischetti et al., 2021) or graded differences in function (Binney et al., 2012; Jackson et al., 2018), including social and semantic function (Olson et al. 2013; Binney et al. 2016). Therefore, under what might be called a *‘dual ATL hub account’*, social conceptual knowledge could be processed within a distinct location to more general conceptual information (Zahn et al. 2007, 2009). Indeed, while social tasks and semantic judgements on social words activate the dorsolateral ATL (the anterior superior and middle temporal gyri; Zahn et al. 2007; Ross and Olson 2010; Binney et al. 2016; Mellem et al. 2016; Lin, Wang, et al. 2018; Wang et al. 2019; Arioli et al. 2020), the general semantics literature, including data from patients and studies using ATL-optimised fMRI (Binney et al. 2010; Mion et al. 2010; Visser et al. 2010; Lambon Ralph et al. 2017), converges on the ventrolateral ATL (including the anterior fusiform and inferior temporal gyri) as the centre-point of a domain-general conceptual hub.

However, two recent studies have demonstrated that using enhanced fMRI techniques greatly affects the patterns of activation observed across the ATL during the processing of social-relevant stimuli and leads to different conclusions (Binney et al. 2016; Rice et al. 2018). In conventional approaches to acquiring fMRI, susceptibility artefacts cause signal loss and image distortion around the location of the ventrolateral ATL, which render the technique effectively blind to activation in this region (Devlin et al. 2000). Spin-echo, and dual-echo echo-planar fMRI, as well as post-acquisition distortion correction techniques, can be used to recover this signal (Embleton et al. 2010; Halai et al. 2014) in which case it becomes clear that the ventrolateral ATL activates strongly during semantic judgements made on both social and non-social stimuli (Hoffman et al. 2015; Binney et al. 2016; Rice et al. 2018). These studies confirmed previous reports by observing that the dorsolateral ATL activates selectively to social semantic stimuli but the omni-category response of the ventrolateral ATL was much greater in magnitude (Binney et al. 2016; Rice et al. 2018). We have argued that these observations are consistent with graded differences in semantic function across a unified representational substrate, as opposed to the ATL comprising two distinct systems for processing social and non-social concepts (Binney et al., 2016). According to the graded semantic hub proposal, a large extent of the ATL contributes to semantic processing in the form of a unified representational space, all of which is engaged by the encoding and retrieval of concepts, and by concepts of any kind. The centre of this space exists over the ventrolateral ATL and its engagement during semantic processing is invariant to, for example, idiosyncratic task features, including the modality through which concepts are accessed. Towards the edges of this space, however, there are gradual shifts in semantic function such that it becomes relatively more specialised for encoding certain types of semantic features (Binney et al. 2012; Rice et al. 2015; Bajada et al., 2019). For example, features that are primarily experienced through certain sensorimotor modalities (for a computational exploration of this general hypothesis, see Plaut 2002). Along these lines, the socialness of a concept might reflect semantic features that are uniquely experienced through certain processing streams like those involved processing affect. The dorsolateral ATL in particular may be sensitive to the socialness of concepts because of strong connectivity to medial temporal limbic and frontal limbic regions (via the uncinate fasciculus; Binney et al. 2012; Bajada et al. 2016; Papinutto et al. 2016). By extension of this proposal, we have argued that activation of ATL subregions in service of social cognitive tasks reflects engagement of a domain-general semantic system and that this is centred upon the ventrolateral ATL (Binney and Ramsey 2020).

The conclusions that can be drawn from those two particular studies regarding the ATL’s role in social cognition are limited. This is because they used tasks where the demands are primarily semantic in nature and the social relevance of the stimuli may have only been a secondary feature. As such, it remains an open question whether social tasks typically employed in the social neuroscience literature activate the ventrolateral ATL hub. The present study tackles exactly that issue, with a specific focus on mental state attribution or ‘theory of mind’ tasks. We chose this focus because theory of mind (ToM) abilities are considered central to the construct of social cognition; they are considered as fundamental to successful social interactions, as they enable us to describe, explain and predict behaviour (Frith and Frith 2005; Brüne and Brüne-Cohrs 2006; Apperly 2012; van Hoeck et al. 2014; Heleven and van Overwalle 2018). Neuroimaging studies reliably implicate the right temporo-parietal junction, medial prefrontal cortex and precuneus as part of a core network for ToM (Saxe and Kanwisher 2003a; Saxe and Wexler 2005; Saxe 2006; Scholz et al. 2009; Young et al. 2010; Dodell-Feder et al. 2011), whereas the role of the ATL is less clear and appears to be characterised as ancillary by some accounts (van Overwalle 2009; Schurz et al. 2014; Molenberghs et al. 2016). It is possible that a central role of the ventrolateral ATL has gone unnoticed because fMRI studies of ToM typically do not account for technical constraints around this region.

We set out to address two key unresolved questions. First, we aimed to determine whether and to what degree different parts of the ATL are activated by established theory of mind tasks. This necessitated two key design elements: (i) the use of dual-echo fMRI and distortion correction to ensure full coverage of the bilateral ATL; and (ii) the use of multiple theory of mind tasks. This second design feature was important because showing common activation across different theory of minds tasks with a variety of stimuli means that we can more confidently assess whether activation can be attributed to ToM ability itself rather than being the result of task demands or idiosyncratic features of the stimuli (Ross and Olson 2010). In fMRI designs, the most commonly used theory of mind tasks include social vignettes, cartoons, and animations that are intended to evoke the attribution of intentions. We used animations as our primary task because they do not directly involve lexical-semantic processing which has been shown to activate the ventrolateral ATL (e. g. Devlin et al. 2000; Spitsyna et al. 2006; Binney et al. 2010; Visser, Embleton, et al. 2010) and would confound our inferences. Moreover, they lend themselves to the creation of a comparable non-social (and non-semantic) control or baseline activation task which is important for controlling for attentional and executive demands, as well as perceptual stimulation. We also acquired data during a False Belief task (Dodell-Feder et al. 2011) and a free-viewing animated film (Jacoby et al. 2016) as these are established paradigms for localising the ‘mentalising’ or theory of mind network.

Second, we set out to directly assess overlap of ToM related activation with activation evoked by semantic decisions made upon nonverbal, non-social stimuli. Overlap, and particularly overlap in the ventrolateral aspect, would support the hypothesis that activation of the ATL during social tasks reflects the retrieval of semantic knowledge representations (Zahn et al. 2007; Olson et al. 2013; Binney and Ramsey 2020). Specifically, we chose the picture version of the Camel and Cactus task (CCT), which is an established means to engage and measure semantic processing, and has been previously used in neuropsychological, functional imaging and brain stimulation studies (Bozeat et al. 2000; Jefferies and Lambon Ralph 2006; Hoffman et al. 2012; Visser et al. 2012). Further, via these means we were also able to directly test three different accounts of the ATL, amongst a single cohort of participants, as follows:

1. The *Social Knowledge Hypothesis/Social Information Processing Account*: according to these accounts, the ATL is part of a domain-specific network involved in processing social information (Simmons et al. 2010), and the dorsolateral ATL is selectively involved in processing concepts with social-emotional content (Olson et al. 2013). Proponents of this account argue against a domain-general role of the ATL in semantic representation (Simmons et al. 2010; Olson et al. 2013). If this is correct, then the ATL would activate for ToM tasks but not during the CCT.
2. The *Dual ATL Hub account*: if a dual hub account is correct then the ToM tasks would exclusively activate the dorsolateral/polar ATL and not the ventrolateral aspect, and the CCT would only activate the ventrolateral ATL.
3. The *Graded ATL Semantic Hub Hypothesis*: if the third account, in which the ventral ATL is the centre point of a domain-general hub for both social and non-social semantic processes (Binney et al. 2016), is correct, then the greatest degree of overlap between the ToM tasks and the CCT will be within the ventrolateral portion. We might also observe dorsolateral activation that is more selective to social stimuli.

## Materials and Methods

### Data Availability statement

Following open science initiatives (Munafò et al. 2017), behavioural and neuroimaging data are openly available on the Open Science Framework project page (https://osf.io/v2gt5/).

### Design Considerations

To ensure that the imaging protocol was sensitive to changes in activation across all parts of the ATL, we used a dual-echo gradient-echo echo-planar imaging (EPI) fMRI sequence that is optimised to detect blood oxygen level-dependent (BOLD) signal in areas of the brain that are usually prone to magnetic susceptibility-induced signal loss (Halai et al. 2014, 2015). Dual-echo sequences like the one used here have been shown to have greater ability to detect inferior temporal lobe activation compared to single-echo GE-EPI sequences with a conventional echo time around 30 ms. (Embleton et al. 2010; Halai et al. 2014). Further, to alleviate the impact of geometric distortions and mislocalisations of fMRI signal also caused by magnetic susceptibility artefacts, the dual-echo sequence was combined with a post-acquisition k-space spatial correction. Our choice of distortion correction procedure was based on evidence that that it outperforms more standard corrections that use phase-encoded field maps (Embleton et al. 2010). We also made other methodological decisions such as acquiring with a left-to-right phase-encoding direction that contributes to better quality of data in the inferior ATL (Embleton et al. 2010). To demonstrate the effectiveness of these measures, we followed the procedure of previous reports investigating the ATLs (Simmons and Martin 2009; Hoffman et al. 2015) and calculated an average map of temporal signal to noise ratio (tSNR) (see Figure M1). The tSNR map suggests a good signal quality even in the in most ventral proportions of the ATL.

Secondly, we adapted stimuli created by Walbrin et al. (2018) to fashion a two alternative force choice (AFC) task involving explicit judgements about the intention of moving shapes (are they behaving in ‘friendly’ or ‘unfriendly’ way?) that are seemingly engaged in goal-directed action. Our task is broadly inspired by the widely used Frith-Happé animations for probing ToM processes (see e.g., Abell et al. 2000; Castelli 2002; Kana et al. 2009; Das et al. 2012; Hennion et al. 2016; Synn et al. 2018; Bliksted et al. 2019) but it was designed to be closer to that used in a key study by Ross and Olsen (2010; also see Shultz et al., 2003). We specifically chose Walbrin and colleagues’ stimuli because they offer a higher number of unique trials (impacting sensitivity/power) than other similar stimuli sets, and they are visually well-controlled to minimize the contribution of low-level visual information to brain responses (i.e., they are comprised of visually diverse interactive scenarios that are well-matched for overall motion energy). To control for attentional and executive demands involved in the main task, we reconfigured these stimuli to further create a well-matched perceptual judgment task. See below for further detail regarding the experimental and control task.

### Participants

Thirty-one healthy native English speakers took part in the experiment. All participants had normal or corrected-to-normal vision, no history of neurological and psychiatric conditions and were right-handed as established by the Edinburgh Handedness Inventory (Oldfield 1971). Participants provided written informed consent, and the study was approved by the local research ethics review committee. Seven participants were excluded because of inadequate task performance (under 70% accuracy) on any one of the social interaction tasks (N=3), or because of failed distortion correction and therefore insufficient data quality (N=4). The final analysed sample comprised of twenty-four participants (12 females, *M*_age_= 22.21, *SD*_age_= 2.13).

### Experimental Stimuli and Tasks

#### Theory of Mind (ToM)

A total of 126 unique video stimuli designed by Walbrin et al. (2018) were used for the main interaction judgement theory of mind (IJ-ToM) task and its corresponding control task. The IJ-ToM stimuli (N = 63) featured two self-propelled circles representing animate agents that were intentionally interacting and doing so in a co-operative manner in half the trials and a competitive manner in the other half (see Supplementary Figure M2). In Walbrin and colleagues’ original stimulus pool (N=256) half of the scenarios concluded with successful goal outcome (e.g., successfully opening a closed door) and half with unsuccessful goal outcomes. Here we only used a subset of the former. In the IJ-ToM task, participants were instructed to make explicit inferential judgements, via a key press, as to whether the agents’ actions towards one another were friendly or unfriendly (unlike the original study that sought to minimize the contribution of ToM judgements and employed a perceptual response task). The associated control task used ‘scrambled’ versions of the interaction stimuli (N = 63) that preserved many of the visual properties but featured altered motion paths such that the shapes did not appear to be intentionally interacting with each other or their environment (see Walbrin et al. 2018). For the present study, these control stimuli were adjusted such that in fifty per cent of trials the speed of motion of one of the two shapes was slower than that of the other. This was done by slowing the frame rate of one of the animation elements (i.e., one of the circles) from 24 to 18 frames per second and removing frames from either the beginning (50%) or end of the sequence to maintain the original duration (6 secs). Moreover, we ensured that the more slowly moving object appeared an equal number of times at each relative position on the screen (e.g., left versus right). Participants responded to these stimuli via key press and indicated whether they believed the circles were moving at same or different speeds. Following some initial pilot behavioural testing, the duration of all 126 videos was shortened from 6 to 3 sec to increase task difficulty/eliminate idle time.

We also acquired data with two widely used functional localisers for the putative ToM network, namely the False Belief (FB) paradigm (Dodell-Feder et al. 2011) and a more recently validated free-viewing movie paradigm (MOV) (Jacoby et al. 2016). The former is a verbal paradigm which is comprised of two sets of 10 text-based vignettes each of which are presented on screen and followed by true / false questions. One of these sets requires the participant to make inferences about a character’s internal beliefs, and this is contrasted against descriptions of facts about physical events. The MOV paradigm involves passive viewing of a commercial animated film and contrasts BOLD responses to events in which characters are involved in ToM against those in which characters experience physical pain (see Jacoby et al. (2016) for more detail).

#### Non-Verbal Semantic Association

Participants also completed a non-verbal version of an established neuropsychological assessment of semantic associative knowledge known as the Camel and Cactus task (CCT; (Bozeat et al. 2000)). This task has been used to engage the semantic network in prior fMRI studies (Visser et al. 2012; Rice et al. 2018). The version used in the present study consisted of 36 trials that contained pictorial stimuli and required participants to make semantic associations between a probe object (e.g., a camel) and a target object (e.g., a cactus) that was presented alongside a foil from the same semantic category (e.g., a rose). The CCT was contrasted against a perceptual control task (36 trials) that consisted of scrambled versions of the CCT pictures and required participants to identify which of two choice pictures was visually identical to a probe (see more detail in Visser et al. (2012)).

#### Experimental Procedure

Participants underwent all testing within a single session lasting approximately one hour. Each individual completed three runs of the IJ-ToM procedure reported below, followed by one run of the CCT procedure, two runs of the FB localiser and one run of the MOV localiser. The IJ-ToM task, the CCT and FB task and the corresponding control tasks were presented via E-prime (Psychology Software Tools, 2017) and the MOV task via Psychtoolbox (Brainard 1997; Pelli 1997) software. Behavioural responses were recorded using an MRI compatible response box.

#### Interaction Judgement ToM Task

The IJ-ToM task and the speed judgement task were paired within a run using an A-rest-B-rest box car block design. Each run contained six blocks per task and three trials per block (18 trials per run per task). There were an equal number of trial types (e.g., *cooperative* versus *competitive*) randomly distributed across blocks within a given run. Both types of active blocks were 17.25 secs long and they were separated by blocks of passive fixation lasting 12 secs each. Each trial began with a fixation cross (duration = 500ms) which was followed by the target animation (3000ms) and finished with a response cue (three question marks; 2000ms). A blank screen occupied an inter stimulus interval of 250ms. Each run lasted 5 minutes and 51 secs and consisted of unique sets of animations. The order in which these runs were completed was counterbalanced across participants. Participants also completed three practice blocks for each of the two tasks before the main runs began.

#### Camel and Cactus Task

The CCT and the corresponding perceptual identity matching control task were alternated within a single run using a blocked design. There were 9 blocks per task, each consisting of four trials (totalling 36 trials per task) and lasting 20 secs. A trial began with a fixation cross (500ms) followed by a stimulus triad (4500ms). Participants responded via key press while the probe and choice items were on screen. Active blocks were separated by brief rest blocks lasting 4000ms and, overall, the run lasted for 7 mins and 12 secs.

#### False Belief and animated movie localisers

Each run of the false belief localiser lasted 4 minutes and 32 seconds and consisted of 10 trials of belief vignettes and 10 trials of the fact vignettes. Finally, the passive MOV scanning run lasted 5 mins and 59 seconds including a fixation period of 10 secs prior to the beginning of the movie. Further details regarding these paradigms are reported by Jacoby et al. (2016).

### Imaging Acquisition

All imaging was performed on a 3T Phillips Achieva MRI scanner with a 32-element SENSE head coil using a 2.5 sense factor for image acquisition. The parameters of the dual-echo gradient-echo EPI fMRI sequence were the following: 31 axial slices covering the whole brain and obtained in an ascending sequential order with a first echo time (TE) = 12ms and second TE = 35ms, repetition time (TR) = 2000ms, flip angle = 85°, FOV (mm) = 240 × 240 × 124, slice thickness = 4 mm, no interslice gap, reconstructed voxel size (mm) = 2.5 × 2.5 and reconstruction matrix = 96 × 96. Prior to image acquisition for each run, we acquired five dummy scans to allow the initial magnetisation to stabilise. This was followed by acquiring 177 volumes for each IJ-ToM task run, 218 volumes for the CCT task run, 138 volumes for each FB task run and 180 volumes for the MOV task run. Adhering to the distortion-correction method, we acquired these functional runs with a single direction k space traversal in the left-right phase-encoding direction. We also acquired a short EPI “pre-scan” with the participants at rest. The parameters of the pre-scan matched the functional scans except that it included interleaved dual direction k space traversals. This gave 10 pairs of images with opposing direction distortions (10 left-right and 10 right-left) which were to be used in the distortion correction procedure described below. To check the quality of the distortion corrected images, we obtained a high resolution T2-weighted scan consisting of 36 slices covering the whole brain, with TR= 17ms, TE= 89ms; reconstructed voxel size (mm) = 0.45 × 0.45 × 4; reconstruction matrix= 512 x512. Additionally, we used a T1-weighted 3D imaging sequence to acquire an anatomical scan, consisting of 175 slices covering the whole brain, for use in spatial normalisation procedures. The parameters of this scan were as follows: P reduction (RL) SENSE factor of 2 and S reduction (FH) SENSE factor of 1, TR = 18ms, TE = 3.4ms, 8° flip angle, reconstructed voxel size (mm) = 0.94 × 0.94 × 1.00 and reconstruction matrix = 240 × 240.

### Data Analysis

#### Behavioural Data

Incorrectly answered trials, missed trials and trials with response latencies that were two standard deviations above or below the participant’s task mean were excluded from analyses of behavioural data. Task performance was assessed in terms of both accuracy and decision times and compared using paired-sample T-tests. Average decision times per block of each task were also calculated so that they could be used as regressors of no interest in fMRI analyses.

#### Distortion Correction and fMRI pre-processing

A spatial remapping correction was computed separately for images acquired at the long and the short echo time, and using a method reported elsewhere (Embleton et al. 2010). This was implemented via in-house MATLAB script (available upon request) as well as SPM12’s (Statistical Parametric Mapping software; Wellcome Trust Centre for Neuroimaging, London, UK) 6-parameter rigid body registration algorithm. Briefly, in the first step, each functional volume was registered to the mean of the 10 pre-scan volumes acquired at the same echo time. Although this initial step was taken primarily as part of the distortion correction procedure, it also functioned to correct the time-series for differences in subject positioning in between functional runs and for minor motion artefacts within a run. Next, one spatial transformation matrix per echo time was calculated from opposingly-distorted pre-scan images. These transformations consisted of the remapping necessary to correct geometric distortion and were applied to each of the main functional volumes. This resulted in two motion-and distortion-corrected time-series per run (one per echo) which were subsequently combined at each timepoint using a simple linear average of image pairs.

All of the remaining pre-processing steps and analyses were carried out using SPM12. Slice-timing correction referenced to the middle slice was performed on the distortion- and motion-corrected images. The T1-weighted anatomical scan was co-registered to a mean of the functional images using a 6-parameter rigid-body transform, and then SPM12’s unified segmentation and normalisation procedure and the DARTEL (diffeomorphic anatomical registration though an exponentiated lie algebra; (Ashburner 2007)) toolbox were used to estimate a spatial transform to register the structural image to Montreal Neurological Institute (MNI) standard stereotaxic space. This transform was subsequently applied to the co-registered functional volumes which were resampled to a 3 × 3x 3 mm voxel size and smoothed with an 8 mm full-width half-maximum Gaussian filter.

#### fMRI Statistical Analysis

Data were analyzed using the general linear model approach (GLM). At the within-subject level, a fixed effect analysis was carried out upon each task pair (e.g., the interaction judgement task and the perceptual control task), incorporating all functional runs within a single GLM. Block onsets and durations were modelled with a boxcar function and convolved with the canonical hemodynamic response function. A high pass filter with a cut off of 128s was also applied. The extracted motion parameters were entered into the model as regressors of no interest. Decision time data were also modelled to account for differences in task difficulty. Due to the block design employed, there was a single value for each epoch of a task which was the average of response times across the trials. These average decision times for each block were mean centred. To avoid false positive activations in the surrounding CSF due to physiological noise, we used an explicit mask restricted to cerebral tissue that was created from tissue segments generated by DARTEL in MNI space and binarised with a 0.4 threshold.

At the level of multi-subject analyses, we first examined activation during the IJ-ToM task at the whole brain level. To complement the whole-brain topographic representation of the data, we used a priori regions of interest (ROI), defined on the basis of independent datasets, to extract and quantify the magnitude of activation within different ATL subregions. This was done using the SPM MarsBar toolbox (Brett et al. 2002). A key aim of this study was also to assess whether parts of the ATL are commonly activated by different types of behavioural paradigm used to localise the putative ToM network. To do this we performed a formal conjunction analysis (Price and Friston 1997; Nichols et al. 2005a). In addition to our interacting geometric shapes paradigm, this included a version of False Belief task (Dodell-Feder et al. 2011) which is comprised of verbal vignettes, and a free-viewing movie paradigm (Jacoby et al. 2016). This analysis was an important step because it would enable us to home in on those regional activations that are a feature of ToM abilities irrespective of the manner in which they are probed. Moreover, if one is to compare ToM tasks that are qualitatively very different from one another, it is possible to attribute the common activations much more convincingly to the particular cognitive process of interest, as opposed to similarities in physical stimulus properties or peripheral elements of the task demands (<u>Friston et al. 1999</u>). Using a further conjunction analysis we also explored overlap between the IJ-ToM task and a nonverbal semantic association task. This enabled us to test the hypothesis that activation of the ATL during social tasks reflects the retrieval of semantic knowledge representations (Zahn et al. 2007; Olson et al. 2013; Binney and Ramsey 2020).

Whole-brain multi-subject random effects analyses were conducted on each of the following contrasts of interest: IJ-TOM task: interaction > speed judgements, interaction judgements > rest, speed judgements > interaction judgements; CCT task: semantic > perceptual judgements; FB task: false belief > false fact judgements; MOV task: mentalizing > pain. One-sample t-tests were performed on all sets of contrast images following application of the same explicit mask as used in the single subject analyses. The resulting statistical maps were assessed for cluster-wise significance using a cluster-defining voxel-height threshold of *p* < .001 uncorrected, and family-wise error (FWE) corrected cluster extent threshold at *p* < .05 (calculated per SPM12 under the random field theory framework; see details regarding smoothness of data, the search volumes and RESELS in **Supplementary Table M1**). Thresholded maps were overlaid on a MNI152 template brain using MRIcroGL (https://www.nitrc.org/projects/mricrogl). We used an AAL atlas implemented in R label4MRI package (https://github.com/yunshiuan/label4MRI) to guide the labelling of peak co-ordinates in the output tables.

Within the ROI analysis, (Brett et al. 2002) two ATL subregions were explored in each hemisphere. A ventrolateral ATL ROI was defined by peak coordinates of activation reported by an independent study of non-verbal semantic processing (Visser et al. 2012) [MNI: +/- 57, -15, -24]. We also examined a polar ATL ROI which was defined on the basis of activation tuned towards socially relevant semantic stimuli as reported by Binney et al. (2016) [MNI: +/-48, 9, -39]. Furthermore, so that we could compare the degree of ATL activation to that of a more established ToM region, we defined a third ROI on the basis of ToM-related TPJ activation reported by Saxe and Kanwisher (2003b) [+/-54, -60, 21]. These sets of coordinates defined a centre of mass for spheres with a radius of 10mm (See **Figure 1 panel B** for an illustration of ROI locations). Per subject, a single summary statistic was calculated to represent activation across all the voxels in an ROI (the mean of the parameter estimates) for the IJ-ToM task relative to the speed judgment control task. One-sample t-tests were then performed to assess group-level significance. To control for multiple comparisons, *p*-values were Bonferroni corrected on the basis of the number of ROIs (multiplied by 6) as implemented in MarsBar. We also conducted planned comparisons between ROIs in each hemisphere, and between hemispheric homologue regions, using paired t-tests. For the conjunction analyses, we used a *p*< .001 uncorrected voxel height threshold to be achieved by each contrast independently prior to conjunction (Price and Friston 1997; Nichols et al. 2005a).

**Figure 1.**
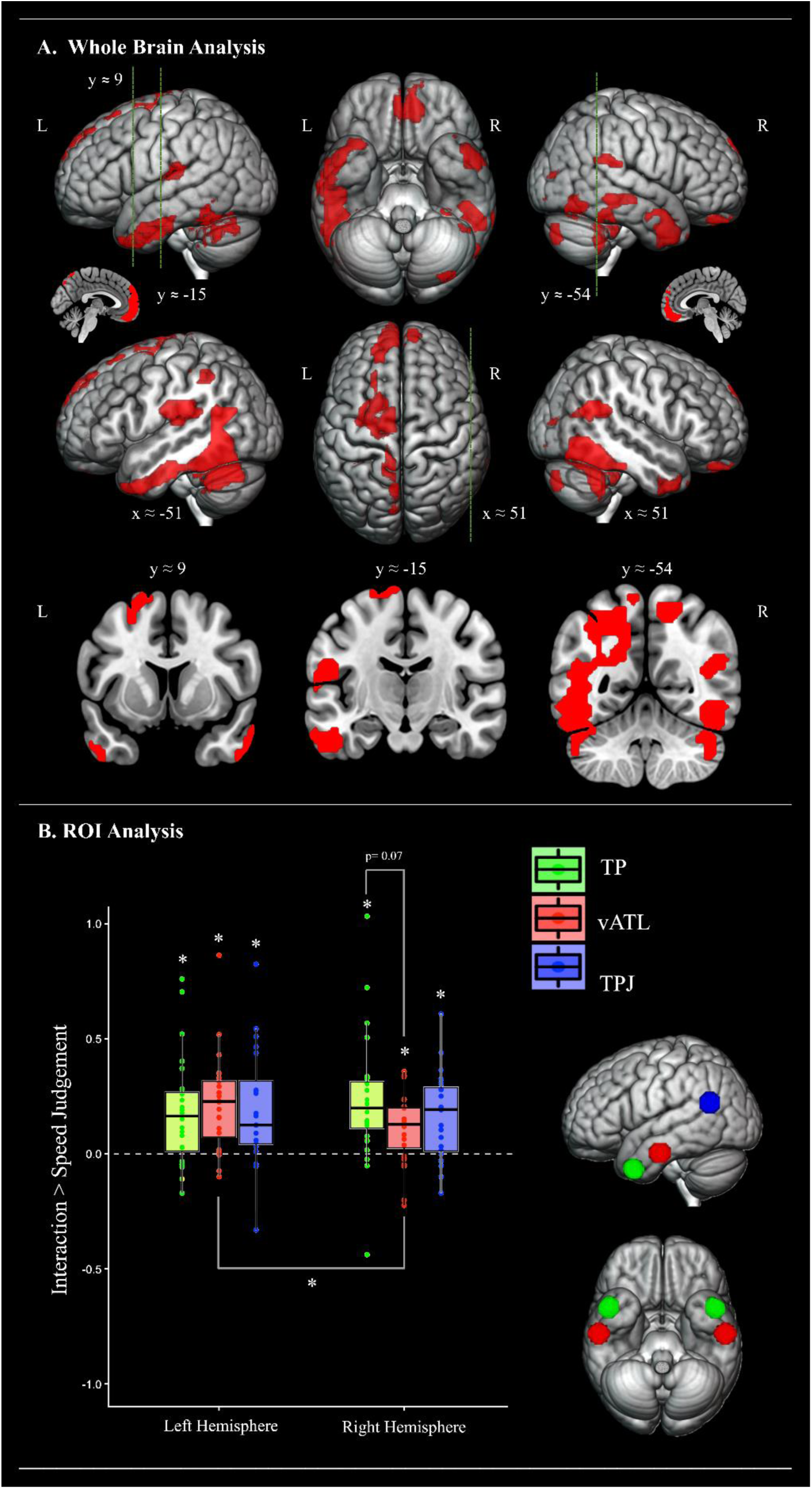
**Panel A.** Cortical regions activated during the main experimental ToM task (the interaction judgement), relative to the speed judgement control task. The statistical map was thresholded with an uncorrected voxel height threshold of p < .001 and a family wise error corrected minimum cluster extent threshold (k= 152) at p < .05. Cross-sections were chosen to display the location of activation found in key studies investigating ToM processing (Saxe & Kanwisher, 2003; right TPJ [51, -54, 27]), semantic processing of social concepts (Binney, Hoffman & Lambon Ralph, 2016; left TP [-48, 9, -39]) and general semantic processing (Visser, Jefferies, Embleton & Lambon Ralph, 2012; left inferior ATL [-57, -15, -24]). Panel B. Summary of the ROI analyses comparing the magnitude of activation for the interaction judgement ToM task (relative to that during speed judgments control task). An asterisk denotes a significant effect at p < .05 after Bonferroni correction. Numerical p-values are displayed where comparisons yielded a p-value greater than .05 but less than .1. TP = temporal pole, vATL = ventrolateral anterior temporal lobe, TPJ = temporo-parietal junction, L = left, R = right.

## Results

### Behavioural Data

Mean accuracy and decision times for all tasks are displayed in **Table 1**. Performance on the animated interaction friendliness judgement (IJ ToM task) was more accurate than on the speed judgement control task (t (23) = 7.50, *p* < .001, Cohen’s *d* = 1.53), and decision times were also faster (t (23) = -3.08, *p* = .005, *d* = 0.63). Performance during semantic association judgements (CCT task) was less accurate than performance in the perceptual identity matching control task (t (23) = -8.83, *p*< .005, *d* = -1.80), although there was no significant difference in the latency of decision times (t (23) = -0.65, p= .522, d= -0.13). Accuracy across the false belief and false facts judgments (FB task) was comparable (t (23) = 0.77, *p* = .450, *d* = 0.16) although decision times were faster in the false fact task (t (23) = 2.73, *p* = .012, *d* = 0.56).

**Table 1.** Behavioural data

| Task | Accuracy (%) | Decision time (ms) |
| --- | --- | --- |
| Interaction Friendliness Judgement | 96.37 (04.35) | 468.38 (125.44) |
| Perceptual Speed Judgement | 84.88 (07.62) | 524.25 (159.84) |
| Semantic Association Judgement | 79.75 (17.84) | 1491.92 (382.65) |
| Perceptual Identity Matching | 92.59 (19.96) | 1529.76 (460.57) |
| False Belief Judgement | 70.42 (19.67) | 2779.56(385.47) |
| False Fact Judgement | 67.92 (15.87) | 2560.31(354.64) |

*Standard deviations stated in parentheses*

### Activation During a Social Attribution Task Given Full Temporal Lobe Coverage

A whole brain univariate analysis contrasting social interaction friendliness judgments with the matched speed judgement task revealed robust bilateral ATL activation that was centred over the ventrolateral aspects in both hemispheres (see **Figure 1, panel A** and **Table 2**). In the left hemisphere, this extended from the ventrolateral temporopolar cortex (BA38), along the inferior middle temporal gyrus and inferior temporal gyrus (ITG), to approximately halfway along the temporal lobe (*y* ≈ -17). This included a maxima that is notably similar in location (MNI coordinates *x* = -54, *y* = 6, *z* = -39) to that identified in association with processing of abstract social concepts (relative to matched abstract non-social concepts; *x* = - 54, *y* = 9, *z* = -33 and animal function concepts; x= -48, y= 9, z= -39) by Binney et al. (2016). The same cluster also extended more posteriorly upon the basal surface and along the fusiform/lingual and posterior inferior temporal gyri. It also traversed up into the parietal lobe and the intraparietal sulcus. In the right hemisphere, ATL activation also covered much of the ventrolateral surface (particularly the polar cortex and the anterior-most portion of the middle temporal gyrus (MTG) but extended less posteriorly (to y ≈ -11) than it did in the left.

**Table 2.** Significant activation clusters in the social interaction judgement > speed judgement contrast (p < .05, FWE-corrected, corresponding to an extent threshold of k = 152 following a cluster-defining threshold of p< .001, uncorrected)

| Cluster Name and Location of Maxima | Cluster Extent (voxels) | Peak (Z) | MNI Coordinates (mm) |  |  |
| --- | --- | --- | --- | --- | --- |
|  |  |  | x | y | z |
| **L temporal – parietal – occipital** | **2242** |  |  |  |  |
| anterior ITG / sulcus |  | 5.72 | -57 | -6 | -30 |
| anterior ITG |  | 5.26 | -54 | 6 | -39 |
| posterior ITG |  | 5.07 | -51 | -51 | -24 |
| precuneus |  | 4.78 | -12 | -60 | 42 |
| posterior ITG |  | 4.76 | -45 | -60 | -9 |
| inferior parietal lobule |  | 4.68 | -39 | -51 | 45 |
| precuneus |  | 4.49 | -9 | -75 | 48 |
| middle/anterior ITG |  | 4.42 | -48 | -21 | -27 |
| posterior MTG |  | 4.23 | -48 | -54 | 3 |
| cerebellum |  | 4.21 | -51 | -51 | -36 |
| posterior MTG |  | 4.10 | -39 | -60 | 15 |
| anterior MTG/TP |  | 4.04 | -42 | 18 | -42 |
| **R temporal – parietal – occipital** | **1708** |  |  |  |  |
| inferior occipital gyrus |  | 5.52 | 48 | -63 | -12 |
| occipital pole |  | 5.01 | 30 | -90 | -6 |
| middle occipital gyrus |  | 4.74 | 33 | -75 | 3 |
| posterior MTG |  | 4.39 | 63 | -36 | -9 |
| cerebellum |  | 4.35 | 36 | -42 | -33 |
| cerebellum |  | 4.29 | 27 | -78 | -42 |
| cerebellum |  | 4.28 | 36 | -84 | -39 |
| posterior STG/TPJ |  | 4.26 | 57 | -39 | 21 |
| cerebellum |  | 4.24 | 42 | -48 | -30 |
| posterior STS/TPJ |  | 4.22 | 42 | -60 | 18 |
| cerebellum |  | 4.21 | 45 | -51 | -42 |
| middle occipital gyrus |  | 4.05 | 39 | -81 | 12 |
| **Bilateral frontal** | **1017** |  |  |  |  |
| L anterior SFG |  | 5.46 | -3 | 63 | 21 |
| R anterior orbital gyrus |  | 5.30 | 9 | 45 | -24 |
| R anterior gyrus rectus |  | 5.21 | 6 | 51 | -18 |
| L middle SFG |  | 5.08 | -12 | 60 | 27 |
| L middle SFG |  | 4.93 | -12 | 57 | 36 |
| R middle SFG |  | 4.81 | 3 | 60 | 30 |
| R anterior mPFC |  | 4.77 | 9 | 63 | -9 |
| R anterior mPFC |  | 4.77 | 3 | 57 | -6 |
| R anterior mPFC |  | 3.45 | 3 | 60 | 6 |
| **L temporal-parietal** | **362** |  |  |  |  |
| superior parietal lobule |  | 4.65 | -54 | -27 | 18 |
| middle STG |  | 3.60 | -63 | -15 | 9 |
| **R anterior temporal** | **160** |  |  |  |  |
| anterior MTG |  | 4.57 | 60 | 6 | -33 |
| anterior ITG |  | 4.54 | 51 | 12 | -45 |
| anterior MTG/TP |  | 3.97 | 51 | 18 | -39 |
| anterior MTG/TP |  | 3.54 | 45 | 24 | -39 |
| anterior MTG |  | 3.37 | 60 | -9 | -24 |
| **L dorsal frontal** | **410** |  |  |  |  |
| middle superior frontal sulcus |  | 4.38 | -24 | 3 | 60 |
| posterior superior frontal sulcus |  | 4.29 | -24 | 0 | 48 |
| posterior SFG |  | 4.15 | -12 | -12 | 78 |
| posterior superior frontal sulcus |  | 4.01 | -30 | -6 | 63 |
| posterior SFG |  | 3.30 | -21 | 27 | 60 |
| **R parietal** | **152** |  |  |  |  |
| superior postcentral gyrus |  | 4.29 | 24 | -42 | 57 |
| superior parietal lobule |  | 4.09 | 18 | -57 | 60 |
| precuneus |  | 3.32 | 9 | -63 | 54 |

*The table shows up to 12 local maxima per cluster more than 8.0 mm apart. L= left; R= right; ITG = inferior temporal gyrus; TP= temporal pole; MTG= middle temporal gyrus; TPJ= temporo-parietal junction; AG= angular gyrus; SFG = superior frontal gyrus; mPFC= medial frontal cortex; STS= superior temporal sulcus*

Outside of the ATL, and as expected, this contrast also revealed activation amongst key nodes of the putative ToM network, including the temporoparietal junction (TPJ), the medial prefrontal cortex (mPFC) and the precuneus (Frith and Frith 2003, 2006; Saxe and Kanwisher 2003b; van Overwalle 2009; Jacoby et al. 2016). TPJ activation was observed in both hemispheres at the position of the posterior superior temporal sulcus/gyrus (STS/STG) and, in the right hemisphere, it extended to more posterior regions (at *y* ≈ -54) that are frequently emphasized in landmark studies (Saxe and Kanwisher 2003b; Saxe and Powell 2006) and large scale meta-analyses (Schurz et al. 2014; Molenberghs et al. 2016) of the theory of mind network. Further activation was revealed in the left posterior MTG, the left insula, and bilateral temporooccipital and cerebellar regions.

Activation during the social interaction friendliness judgments was also contrasted with passive fixation/rest. There was notably little activation in the ATLs, except for a small cluster in the left superior temporal pole (see **Supplementary Figure 1** and **Supplementary Table R1**). This is consistent with the idea that there may be automatic semantic activation (e.g., mind-wandering, episodic recall and socially oriented thoughts) during periods of passive fixation, and it demonstrates the importance of using active baseline tasks for detecting ATL activation which has been highlighted in prior meta-analyses and empirical investigations (Binder et al. 1999, 2009; Visser, Jefferies, et al. 2010; Andrews-Hanna et al. 2014).

Outside of this region there was robust bilateral fronto-parietal activation including of the bilateral TPJ and the ventrolateral prefrontal cortex, and activation of the mPFC, the precuneus, and temporooccipital and cerebellar regions. The contrast revealing greater activation for the speed judgment relative to the social attribution task is reported in **Supplementary Figure 2** and **Supplementary Table R2** and revealed the right middle frontal gyrus and a number of midline structures.

We used an a priori ROI-based approach to compare the magnitude of regional responses to the social attribution task both within each hemisphere and between hemispheric homologues. We focused upon two key ATL subregions, the temporopolar cortex and the posteriorly adjacent ventrolateral surface, as well as temporoparietal cortex (i.e., the TPJ) frequently implicated in theory of mind. The positions of these ROIs and the results are displayed in **Figure 1, panel B**. Bonferroni-corrected one-sample T-tests revealed significant activation during social interaction judgements in the left vATL (t (24) = 5.09, Cohen’s d= 1.04), temporal pole (t (24) = 3.77, Cohen’s d= .77) and TPJ (t (24) = .20, Cohen’s d= .04) and also the right vATL (t (24) = 3.57, Cohen’s d= .73), temporal pole (t (24) = 4.03, Cohen’s d= .82) and TPJ (t (24) = .17, Cohen’s d= .04) (all *p* < .005). Numerically speaking, across all the ROIs, the left vATL revealed the largest effect size, and the TPJ showed the weakest effects. Planned statistical comparisons (**see Supplementary Table R3**) confirmed greater activation in the left as compared to the right vATL (t (24) = 2.45, *p* = .02, Cohen’s *d* =.50). There were no other significant pairwise differences.

### Common Activation of the ATL Across Three Different ToM Paradigms

In the subsequent analysis, we aimed to map out subregions of the bilateral ATL in which there is overlapping activation between some of the different types of behavioural paradigm used to localise the putative ToM network (Dodell-Feder et al. 2011; Jacoby et al. 2016). The results of independent whole-brain analyses contrasting two further ToM tasks (the False Belief task and the free-viewing movie paradigm) with their respective control tasks are reported in **Supplementary Figures 3** and **4 and Supplementary Tables R4 and R5**. We formally assessed activation overlap between the three ToM tasks using a conjunction analysis performed across the whole brain (Nichols et al. 2005b). For complete visualisation of the results and to capture the full extent of both the overlap and divergence in the topography of activation, the three whole brain activation maps are overlaid on each other in **Figure 2 panel A**, whereas a map limited to the formal statistical conjunction can be found in **Supplementary Figure 5** and **Supplementary Table R6**. Regarding ATL activation, the conjunction analysis revealed three-way overlap between the ToM tasks exclusively within the left ventrolateral ATL. This extended over the anterior ITG and MTG from about y ≈ -7 to y ≈ 9 and is also strikingly similar to ATL regions reported as activated by social concepts by (Binney, et al. 2016). As would be expected from prior literature, 3-way overlap was also observed in the mPFC and bilateral TPJ.

**Figure 2.**
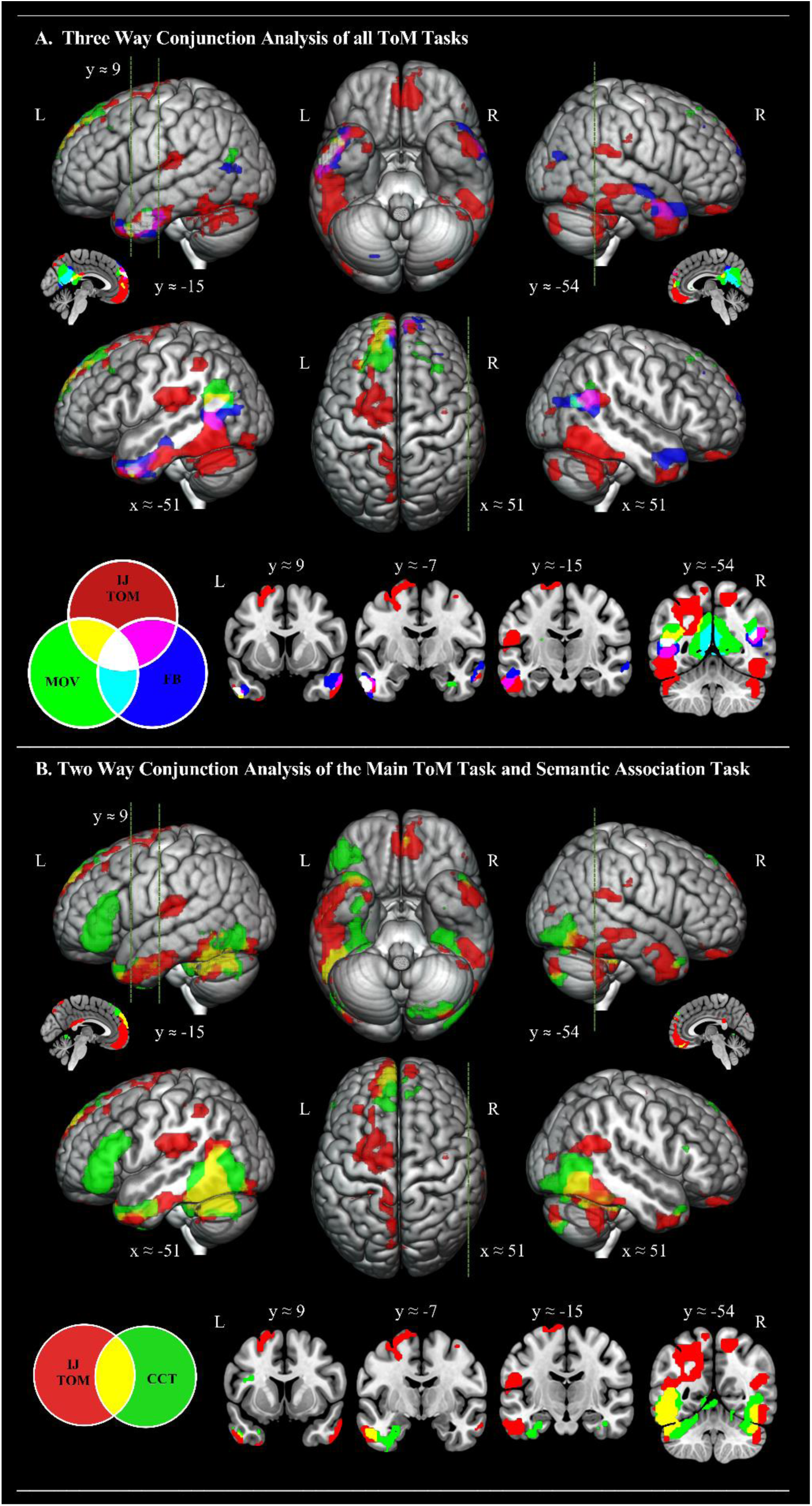
**Panel A.** Topological overlap of cortical regions activated by the interaction judgement > speed judgement contrast, the false belief story > photograph contrast, and the mentalising > pain contrast from the free-viewing movie localiser. Each of the three statistical maps were independently thresholded with an uncorrected voxel height threshold of p < .001 and then overlaid within MRICron using additive colour blending. White patches indicate three-way overlap between all three ToM contrasts. Panel B. Topological overlap of cortical regions activated by the interaction judgement > speed judgement contrast (red), and the nonverbal semantic association (Camel and Cactus task) > perceptual judgement contrast (green). The two statistical maps were independently thresholded with an uncorrected voxel height threshold of p < .001 and then overlaid within MRICron using additive colour blending. Yellow patches indicate overlap between theory of mind and general semantic processing. Cross-sections were chosen to display the location of activation found in key studies investigating ToM processing (Saxe & Kanwisher, 2003; right TPJ [51, -54, 27]), semantic processing of social concepts (Binney, Hoffman & Lambon Ralph, 2016; left TP [-48, 9, -39]) and general semantic processing (Visser, Jefferies, Embleton & Lambon Ralph, 2012; left inferior ATL [-57, -15, -24]), as well one further key area of 3-way overlap (y = -7).

### ATL Activation Common to both ToM and General Semantic Processing

Finally, we performed a conjunction analysis aimed at identifying any potential overlap between ATL regions engaged by theory of mind tasks and those engaged by general semantic processing. The same sample of participants and the same ATL-optimised dual-echo imaging sequence were used to acquire fMRI data while individuals completed a nonverbal semantic association task. The result of an independent whole-brain analysis contrasting this task with a matched control task is reported in **Supplementary Figure 6** and **Supplementary Tables R7**. This contrast was entered into a whole brain conjunction analysis along with the interacting geometric shapes paradigm. The full extent of overlap and divergence between ToM activation and general semantic activation is displayed in **Figure 2 panel B**, while the results of the formal statistical conjunction are found in **Supplementary Figure 7** and **Supplementary Table R8**. Both theory of mind and general semantics activated the left ventrolateral ATL. Specifically, there was a cluster of 114 commonly activated voxels in the left ventral ATL with the activation starting to converge at y ≈ -15, showing the most robust overlap at y ≈ -7, and still overlapping in inferior polar regions at y ≈ 9. There was a further common ATL activation (extent = 32 voxels) within the left medial temporal pole. On the basis of this analysis, the right ATL appeared only to be activated by the IJ-ToM task. Outside of the ATL region, there was also overlap in the left pMTG and TPJ region, as well as the left mPFC and bilateral inferior temporo-occipital regions.

## Discussion

The present study was aimed at evaluating alternative accounts of the role of the anterior temporal lobes (ATL) in social cognition. According to the *social knowledge hypothesis* and the *Social Information Processing Account*, this region serves a domain-specific role in processing socially-relevant semantic information, and its proponents have argued against any domain-general role in semantic processing (Simmons et al. 2010; Olson et al. 2013). Further, this proposal particularly pinpoints the dorsolateral subregions of this relatively large and structurally heterogenous area (Zahn et al. 2007; Ross and Olson 2010). Another hypothesis, which we refer to as the ‘*dual hub’* account, distinguishes between two separate ATL hubs, one for social semantic and one for general semantic processing. Alternatively, the *graded semantic hub* hypothesis, holds that the ATL is a unified domain-general conceptual hub involved in the representation of all manner of conceptual-level knowledge (Binney, et al. 2016; Lambon Ralph et al. 2017). According to this account, the ventrolateral ATL is a critical centre-point for general semantic knowledge representation. Other ATL sub-regions, including the dorsolateral surface and the poles, are characterised as having connectivity-driven graded variations in semantic function, including a ‘sensitivity’ to information that is perceived primarily within certain sensorimotor modalities and/or has a particular behavioural (e.g., social) relevance (Plaut 2002; Visser and Lambon Ralph 2011; Binney et al. 2012; Binney et al. 2016). The key findings of the present study were as follows:

1. By using distortion-corrected dual-echo fMRI, we were able to confirm, within a whole-brain analysis, that the bilateral ventrolateral ATL is engaged by a nonverbal task of the kind that has frequently been employed to localise the putative theory of mind network. A region of interest analysis revealed that the left ventrolateral ATL was more activated than the right homologue, and both these regions were as robustly activated as the TPJ which is another key node in the theory of mind network.
2. Moreover, the left ventrolateral ATL activation was confirmed as a key feature of theory of mind by the fact that it was activated robustly across three different paradigms employing a range of verbal and nonverbal stimuli.
3. Finally, the left ATL activation associated with theory of mind greatly overlapped with that evoked by semantic association judgements performed on non-social picture stimuli.

Overall, these findings support the hypothesis that the ATL is a domain-general conceptual hub and suggest that its contribution to social cognition is related to the retrieval of a general class of semantic knowledge representations (Binney and Ramsey 2020). These findings are not compatible with the social knowledge hypothesis nor the dual hub hypothesis

### The Functional Contribution of ATL Subregions to Social and Semantic Cognition

A link between certain parts of the ATL (e.g., temporopolar cortex; for a review see Olson et al. 2013) and social cognition has been recognised for well over a century, owed in part to the acclaimed work of Brown and Schafer (1888) and, later, Klüver and Bucy (1937) who performed bilateral ATL resection in non-human primates. These investigations are best known for the profound post-operative changes in social behaviour, including emotional blunting and hypersexuality. However, Klüver and Bucy’s primary aims were to establish whether these bilateral lesions led to high-level perceptual deficits, namely visual and auditory associative agnosias or, as referred to by these authors, ‘psychic blindness’. Indeed, this set of studies detail a broad symptom complex that was chiefly characterised by a failure to generate the meaning of visual and auditory stimuli. Therefore, it appears that their subjects were exhibiting multimodal semantic deficits that might explain, and not just co-present with, the social-affective disturbances.

In more recent years, the social neurosciences have seen another rise in interest regarding the specific role played by the ATL (for a review see Olson et al. 2013). In particular, there emerged the *social knowledge hypothesis*, which states that this region supports a domain-specific class of semantic knowledge: social concepts (Zahn et al. 2007; Ross and Olson 2010; Simmons et al. 2010). Although this account acknowledges supporting evidence from within comparative and behavioural neurology, it is primarily based on functional neuroimaging data which specifically points to the dorsolateral and polar ATL (also see Zahn et al., 2007).

Another long-standing series of studies have implicated the ATL in more general forms of semantic processing (Lambon Ralph et al. 2017). These include detailed neuropsychological investigations of a disorder known as semantic dementia (SD). The SD syndrome falls within the spectrum of frontotemporal dementia and exhibits relatively focal atrophy and hypometabolism centred on the bilateral anterior temporal lobes (Mummery et al. 2000; Nestor et al. 2006). This is coupled with a progressive, central impairment of semantic memory that is evident in both expressive and receptive semantic tasks, and across all modalities including spoken and written language, object use, picture-based tasks, environmental sound tasks, and in olfaction and taste (Hodges and Patterson 2007; Luzzi et al. 2007; Patterson et al. 2007; Piwnica Worms et al. 2010). Moreover, this human disorder displays striking parallels to the observations of Klüver and Bucy, in that the multimodal semantic deficit is accompanied by a range of socio-affective deficits, which include impaired emotion recognition and empathy, impaired capacity for ToM, and a loss of person-specific knowledge (Edwards-Lee et al. 1997; Binney et al. 2016; Snowden et al. 2018; Ding et al. 2020). This patient evidence is bolstered by a now extensive set of multi-method studies that used electrophysiological recordings, neurostimulation techniques (TMS/tDCS) and/or functional neuroimaging in neurotypical samples (Marinkovic et al. 2003; Pobric et al. 2008; Binney et al. 2010; Chan et al. 2011; Binney and Lambon Ralph 2015; Shimotake et al. 2015) all of which point to a role of the ATL in general semantic processing. However, as compared to the social knowledge hypothesis, this literature has converged upon a different subregion, the ventrolateral ATL, as the critical substrate for semantic knowledge representation. This includes the findings of ATL-optimised fMRI studies and the data from SD which reveals that the ventrolateral ATL is, alongside the temporopolar regions, the most atrophied ATL subregion in this disorder (Galton et al. 2001; Binney et al. 2010; Mion et al. 2010). Moreover, it is noteworthy that Klüver and Bucy (1939) also remarked that the symptoms they observed in non-human primate’s failed to appear after resections limited to the dorsolateral convolutions of the temporal lobe. Nor did they present after severing connections of the temporal lobe to the frontal or to the occipital lobes.

The findings of the present study are most compatible with this second set of observations and implicate the ventrolateral ATL in both social and general semantic processing. To our knowledge, they represent the first firm demonstration using fMRI of ventrolateral ATL activation during the types of social (and more specifically, theory of mind) paradigms that are typically employed in the social neuroscience literature. This ATL subregion is frequently missing from fMRI studies probing theory of mind because of methodological considerations we were able to overcome (see below). The fact that three very different theory of mind paradigms evoked ventrolateral ATL activation suggest that it is a feature of ToM irrespective of the paradigm with which it is probed and therefore that it reflects a core cognitive component of theory of mind. Moreover, the fact that this activation overlapped directly with that evoked by a set of non-social semantic judgements is consistent with the claim that engagement of the ATL by social tasks reflects access to a general class of domain-general conceptual representations (Binney and Ramsey 2020).

Our results complement recent studies that found evidence of a role of the left ventrolateral ATL in accessing abstract social concepts (Binney et al. 2016; Rice et al. 2018) as well as other forms of social conceptual knowledge such as person semantics (Rice et al. 2018). Moreover, this region responds to meaningful stimuli across number of domains including faces, bodies, objects, and linguistic stimuli (Visser et al. 2012; Avidan et al. 2014; Collins and Olson 2014; Harry et al. 2016; Ramot et al. 2019). This, alongside the fact that we were able to directly demonstrate ventrolateral ATL activation common to both nonverbal (the interacting shapes task) and verbal (the false belief vignettes) theory of mind tasks is consistent with the notion that the ventrolateral ATL is a supramodal hub engaged in semantic retrieval irrespective of the sensory, motor or linguistic modality through which concepts are probed (Lambon Ralph et al. 2017).

The dorsolateral ATL subregion previously implicated in the representation of social conceptual knowledge (e.g., Zahn et al. 2007) was notably absent within our main set of contrasts. This was unexpected and the reasons are unclear. It is important to note however that an absence of evidence for activation within this and other ATL subregions does not necessarily preclude the possibility that they also have a role in theory of mind and/or general semantic processing. Instead, it is possible that at the level of stringent thresholding that we employed in this study we have only been able to identify the strongest peaks of activation, or functional epicenters. In this case, it might be that activation in other ATL subregions only becomes detectable at a supra-threshold level when there is increased power from higher levels of sampling or when more finely tuned contrasts are used to evoke greater effects sizes. We did contrast semantic judgments made on social and non-social stimuli in two prior studies, and these revealed a sensitivity of activation to social stimuli in the dorsolateral ATL (Binney et al. 2016; Rice et al. 2018). However, these findings do not support a dual hub account of the ATL in which there are discrete functional subdivisions for difference classes of concept. Instead, they were in alignment with a ‘graded hub’ account in which the whole ATL comprises a single semantic hub but it has graded subspecialisations towards certain types of conceptual information (Plaut 2002; Binney et al. 2012; Rice et al. 2015). This is because the adjacent ventrolateral ATL responded equally to both the social and non-social stimuli, and to a much greater extent than the dorsolateral subregion. According to graded hub hypothesis, the ventrolateral ATL region is the centre-point of the hub and has a modality/domain/category-general semantic function. The sensitivity of the dorsolateral/polar ATL to social stimuli may follow from this subregion’s close proximity to and strong connectivity with the limbic system (via the uncinate fasciculus; Binney et al. 2012; Papinutto et al. 2016; Bajada et al. 2017), and could reflect a specialisation in the assimilation of, for example, emotion-related or interoceptive information into coherent semantic representations (Olson et al. 2007; Vigliocco et al. 2014; Rice et al. 2015).

A clear difference the way in which the ATL was engaged by the semantic judgements and the theory of mind tasks is that the latter appears far more bilateral. Moreover, ToM elicited bilateral ATL activation regardless of the verbal/non-verbal nature of the stimuli. The role of the ATL in semantic cognition is proposed to be bilateral although, again, perhaps with graded specialisations towards processing verbal semantic information in the left hemisphere (Lambon Ralph et al. 2001; Rice et al. 2015). The role of the ATL in social cognition has been ascribed with a right lateralisation within some accounts drawing mainly on neuropsychological patient evidence (Gorno-tempini et al. 2003; Zahn et al. 2009; Irish et al. 2014; Gainotti 2015; Rice et al. 2017; Borghesani et al. 2019) although the fMRI studies reviewed above (Ross and Olson 2010; Binney et al. 2016; Rice et al. 2018) indicate bilateral involvement (also see Rice, Lambon Ralph, et al. 2015; Pobric et al. 2016; Lin et al. 2018; Arioli et al. 2020; Catricalà et al. 2020). Contrary to all these prior findings, the present ROI results are suggestive of a greater response in the left as compared to the right ATL. The reasons for this are unclear, and an interesting aim for future neuroimaging studies is to explore factors (e.g., task; stimulus modality) that could potentially drive differences in the activation of bilateral ATL subregions both in the context of social and general semantic tasks.

### The status of the ATL in neurobiological accounts of social cognition

Animal ablation studies (Brown and Schafer 1888; Klüver and Bucy 1937) and case descriptions of the profound consequences for humans of focal ATL lesions (Terzian and Dalle Ore 1955) and degeneration (e.g. Edwards-Lee et al. 1997) provided some relatively early clues as to the importance of the anterior temporal cortex for socio-affective competences. Nonetheless, the ATL often does not feature prominently within contemporary neurobiological frameworks for understanding social behaviour (Decety and Lamm 2007; Lieberman 2007; Adolphs 2009; van Overwalle 2009; Spunt and Adolphs 2017). It is overshadowed by prefrontal, medial and lateral temporoparietal regions, and seemingly attributed with an ancillary status. This could be due, at least in part, to the predominance of fMRI in the social neurosciences and the fact that this technique is typically blind to activation in a significant proportion of this region (Devlin 2002). Inconsistencies in the presence and location of ATL activation across various social domains, relative to the TPJ for example, could explain a modest appetite for further exploring the region’s contribution.

Here, and in two prior ATL-optimised fMRI studies (Binney et al. 2016; Rice et al. 2018), we have shown that when steps are taken to alleviate the technical limitations of the fMRI technique, robust ATL activations are observed across a variety of social stimuli and social tasks. Activation also occurs in a ventrolateral ATL region that is one of the most affected in patients with both striking semantic and social impairments (Binney et al. 2010; Binney et al. 2016; Kumfor et al. 2016). Moreover, in the present study, we have demonstrated that left ventrolateral ATL activation is at least as robust, in extent and magnitude, as that of another key social region (the TPJ), and at least as consistent across different tasks and stimuli. Overall, we interpret this as initial evidence from neurotypical samples to complement that obtained from patient studies, that the ventrolateral ATL is of equal functional import to social cognition as other key nodes of the ‘social brain’ (such as the TPJ, the mPFC and the precuneus).

Several authors have argued that progress in social neuroscience theory will rapidly accelerate if it embraces established and detailed models from within other more general domains of cognition (Spunt and Adolphs 2017; Amodio 2019; Ramsey and Ward 2020). Taking a similar perspective, we have recently proposed that a unifying feature amongst many forms of social cognitive processing is the retrieval of conceptual knowledge, and that it could be productive to understand social cognition to essentially be an example of semantic cognition (Binney and Ramsey 2020). This would appear a reasonable viewpoint given that social interaction is, at its core, a process of *meaningful* exchange between persons. The main practical implication of this proposal, at least for the present discussion, is that social and semantic cognition rely on the same cognitive and brain mechanisms, and this positions the ventrolateral ATL at the heart of social cognition. According to this framework, other key nodes of the ‘social brain’, including the mPFC and the TPJ, could also serve a domain-general role rather than one that is specialised towards processing social information (van Overwalle 2009; Seghier et al. 2010; Cabeza et al. 2012; Bzdok et al. 2016; Humphreys et al. 2020; Diveica et al. 2021). In summary, we argue that there is a growing need to re-evaluate the relative contribution of all these regions, as well as develop a better understanding of the way they interact in service of social cognition.

### Conclusions and Future Directions

In conclusion, our findings support the claim that the ventrolateral ATL is an important contributor to social cognition and point to a specific role as a domain-general hub for conceptual knowledge representations that help inform our understanding of others and guide our own meaning-driven social behaviours. A key methodological determinant underpinning these findings was the use of a neuroimaging technique that maximises the signal obtained from across the entire ATL region. However, the present study is also limited by its methodology. To a large extent, fMRI remains the predominant mode of investigation in the social neurosciences. However, it cannot be escaped that the inferences it allows are merely correlational and not at all causal. For this reason, the field needs to increasingly turn to patient models such as stroke, temporal lobe epilepsy, and frontotemporal dementia (Kumfor et al. 2017; Rankin 2020, 2021), as well as non-invasive techniques, such as transcranial magnetic stimulation, that can be used to more directly probe the neural architecture of cognition in neurological healthy samples. This will enable us to get a firmer grasp on key questions including those regarding the laterality of function within the ATL and the TPJ, as well as the functional necessity of distinct subregions.

## Acknowledgements

The authors would like to thank Jordan Bryne and Taylor Baumler for their assistance with data collection and recruitment, and Paul Downing and Kami Koldewyn for their comments on a previous version of this manuscript.

## References

Abell F, Happé F, Frith U. 2000. Do triangles play tricks? Attribution of mental states to animated shapes in normal and abnormal development. Cognitive Development. 15:1–16.

Adolphs R. 2009. The Social Brain: Neural Basis of Social Knowledge. Annual Review of Psychology. 60:693–716.

Amodio DM. 2019. Social Cognition 2.0: An Interactive Memory Systems Account. Trends in Cognitive Sciences. 23:21–33.

Andrews-Hanna JR, Smallwood J, Spreng RN. 2014. The default network and self-generated thought: component processes, dynamic control, and clinical relevance. Annals of the New York Academy of Sciences. 1316:29–52.

Apperly IA. 2012. What is “theory of mind”? Concepts, cognitive processes and individual differences. Quarterly Journal of Experimental Psychology. 65:825–839.

Arioli M, Gianelli C, Canessa N. 2020. Neural representation of social concepts: a coordinate-based meta-analysis of fMRI studies. Brain Imaging and Behavior.

Ashburner J. 2007. A fast diffeomorphic image registration algorithm. NeuroImage. 38:95–113.

Avidan G, Tanzer M, Hadj-Bouziane F, Liu N, Ungerleider LG, Behrmann M. 2014. Selective dissociation between core and extended regions of the face processing network in congenital prosopagnosia. Cerebral Cortex. 24:1565–1578.

Bajada CJ, Haroon HA, Azadbakht H, Parker GJM, Lambon Ralph MA, Cloutman LL. 2017. The tract terminations in the temporal lobe: Their location and associated functions. Cortex. 97:277–290.

Binder JR, Desai RH, Graves WW, Conant LL. 2009. Where is the semantic system? A critical review and meta-analysis of 120 functional neuroimaging studies. Cerebral Cortex. 19:2767–2796.

Binder JR, Frost JA, Hammeke TA, Bellgowan PSF, Rao SM, Cox RW. 1999. Conceptual Processing during the Conscious Resting State: A Functional MRI Study. Journal of Cognitive Neuroscience. 11:80–93.

Binney RJ, Embleton K v., Jefferies E, Parker GJMM, Lambon Ralph MA. 2010. The ventral and inferolateral aspects of the anterior temporal lobe are crucial in semantic memory: Evidence from a novel direct comparison of distortion-corrected fMRI, rTMS, and semantic dementia. Cerebral Cortex.

Binney RJ, Henry ML, Babiak M, Pressman PS, Santos-Santos MA, Narvid J, Mandelli ML, Strain PJ, Miller BL, Rankin KP, Rosen HJ, Gorno-Tempini ML. 2016. Reading words and other people: A comparison of exception word, familiar face and affect processing in the left and right temporal variants of primary progressive aphasia. Cortex. 82:147–163.

Binney RJ, Hoffman P, Lambon Ralph MA. 2016. Mapping the Multiple Graded Contributions of the Anterior Temporal Lobe Representational Hub to Abstract and Social Concepts: Evidence from Distortion-corrected fMRI. Cerebral Cortex. 26:4227–4241.

Binney RJ, Lambon Ralph MA. 2015. Using a combination of fMRI and anterior temporal lobe rTMS to measure intrinsic and induced activation changes across the semantic cognition network. Neuropsychologia. 76:170–181.

Binney RJ, Parker GJM, Lambon Ralph MA. 2012. Convergent Connectivity and Graded Specialization in the Rostral Human Temporal Lobe as Revealed by Diffusion-Weighted Imaging Probabilistic Tractography. Journal of Cognitive Neuroscience. 24:1998–2014.

Binney RJ, Ramsey R. 2020. Social Semantics: The role of conceptual knowledge and cognitive control in a neurobiological model of the social brain. Neuroscience and Biobehavioral Reviews.

Bliksted V, Frith C, Videbech P, Fagerlund B, Emborg C, Simonsen A, Roepstorff A, Campbell-Meiklejohn D. 2019. Hyper- and Hypomentalizing in Patients with First-Episode Schizophrenia: fMRI and Behavioral Studies. Schizophrenia Bulletin. 45:377–385.

Borghesani V, Narvid J, Battistella G, Shwe W, Watson C, Binney RJ, Sturm V, Miller Z, Mandelli ML, Miller B, Gorno-Tempini ML. 2019. “Looks familiar, but I do not know who she is”: The role of the anterior right temporal lobe in famous face recognition. Cortex.

Bozeat S, Lambon Ralph MA, Patterson K, Garrard P, Hodges JR. 2000. Non-verbal semantic impairment in semantic dementia. Neuropsychologia. 38:1207–1215.

Brainard DH. 1997. The Psychophysics Toolbox. Spatial Vision. 10:433–436.

Brett Anton J. L. Valabregue R. & Poline J. B. M, Brett, M., Anton, J. L., Valabregue, R., & Poline JB. 2002. Region of interest analysis using the MarsBar toolbox for SPM 99. Neuroimage. 16.

Brown S, Sharpey-Schafer EA. 1888. XI. An investigation into the functions of the occipital and temporal lobes of the monkey’s brain. Philosophical Transactions of the Royal Society of London(B). 303–327.

Brüne M, Brüne-Cohrs U. 2006. Theory of mind-evolution, ontogeny, brain mechanisms and psychopathology. Neuroscience and Biobehavioral Reviews.

Bzdok D, Hartwigsen G, Reid A, Laird AR, Fox PT, Eickhoff SB. 2016. Left inferior parietal lobe engagement in social cognition and language. Neuroscience and Biobehavioral Reviews. 68:319–334.

Cabeza R, Ciaramelli E, Moscovitch M. 2012. Cognitive contributions of the ventral parietal cortex: an integrative theoretical account. Trends in Cognitive Sciences. 16:338–352.

Castelli F. 2002. Autism, Asperger syndrome and brain mechanisms for the attribution of mental states to animated shapes. Brain. 125:1839–1849.

Catricalà E, Conca F, Fertonani A, Miniussi C, Cappa SF. 2020. State-dependent TMS reveals the differential contribution of ATL and IPS to the representation of abstract concepts related to social and quantity knowledge. Cortex. 123:30–41.

Chan AM, Baker JM, Eskandar E, Schomer D, Ulbert I, Marinkovic K, Cash SS, Halgren E. 2011. First-pass selectivity for semantic categories in human anteroventral temporal lobe. Journal of Neuroscience. 31:18119–18129.

Collins JA, Koski JE, Olson IR. 2016. More Than Meets the Eye: The Merging of Perceptual and Conceptual Knowledge in the Anterior Temporal Face Area. Frontiers in Human Neuroscience. 10:189.

Collins JA, Olson IR. 2014. Knowledge is power: How conceptual knowledge transforms visual cognition. Psychonomic Bulletin and Review. 21:843–860.

Das P, Lagopoulos J, Coulston CM, Henderson AF, Malhi GS. 2012. Mentalizing impairment in schizophrenia: A functional MRI study. Schizophrenia Research. 134:158–164.

Decety J, Lamm C. 2007. The role of the right temporoparietal junction in social interaction: How low-level computational processes contribute to meta-cognition. Neuroscientist. 13:580–593.

Devlin J. 2002. Is there an anatomical basis for category-specificity? Semantic memory studies in PET and fMRI. Neuropsychologia. 40:54–75.

Devlin JT, Russell RP, Davis MH, Price CJ, Wilson J, Moss HE, Matthews PM, Tyler LK. 2000. Susceptibility-Induced Loss of Signal: Comparing PET and fMRI on a Semantic Task. NeuroImage. 11:589–600.

Ding H, Jiang X, Member S, Shuai B, Liu AQ, Wang G, Member S. 2020. and Multi-Path Decoding. 29:3520–3533.

Ding S-L, van Hoesen GW, Cassell MD, Poremba A. 2009. Parcellation of human temporal polar cortex: A combined analysis of multiple cytoarchitectonic, chemoarchitectonic, and pathological markers. The Journal of Comparative Neurology. 514:595–623.

Diveica V, Koldweyn K, Binney RJ. 2021. Establishing a Role of the Semantic Control Network in Social Cognitive Processing: A Meta-analysis of Functional Neuroimaging Studies. bioRxiv.

Dodell-Feder D, Koster-Hale J, Bedny M, Saxe R. 2011. FMRI item analysis in a theory of mind task. NeuroImage. 55:705–712.

Edwards-Lee T, Miller BL, Benson DF, Cummings JL, Russell GL, Boone K, Mena I. 1997. The temporal variant of frontotemporal dementia. Brain. 120:1027–1040.

Embleton K v., Haroon HA, Morris DM, Ralph MAL, Parker GJMM. 2010. Distortion correction for diffusion-weighted MRI tractography and fMRI in the temporal lobes. Human Brain Mapping. 31:1570–1587.

Frith C, Frith U. 2005. Theory of mind. Current Biology. 15:R644–R645.

Frith CD, Frith U. 2006. The Neural Basis of Mentalizing. Neuron. 50:531–534.

Frith U, Frith C. 2010. The social brain: allowing humans to boldly go where no other species has been. Philosophical Transactions of the Royal Society B: Biological Sciences. 365:165–176.

Frith U, Frith CD. 2003. Development and neurophysiology of mentalizing. Philosophical Transactions of the Royal Society of London Series B: Biological Sciences. 358:459–473.

Gainotti G. 2015. Is the difference between right and left ATLs due to the distinction between general and social cognition or between verbal and non-verbal representations? Neuroscience and Biobehavioral Reviews. 51:296–312.

Gallagher HL, Frith CD. 2003. Functional imaging of ‘theory of mind. ’ Trends in Cognitive Sciences. 7:77–83.

Galton CJ, Patterson K, Graham K, Lambon-Ralph MA, Williams G, Antoun N, Sahakian BJ, Hodges JR. 2001. Differing patterns of temporal atrophy in Alzheimer’s disease and semantic dementia. Neurology. 57:216–225.

Gorno-tempini ML, Rankin KP, Woolley JD, Rosen HJ, Phengrasamy L, Miller BL. 2003. Right Temporal Atrophy.

Halai AD, Parkes LM, Welbourne SR. 2015. Dual-echo fMRI can detect activations in inferior temporal lobe during intelligible speech comprehension. NeuroImage. 122:214–221.

Halai AD, Welbourne SR, Embleton K, Parkes LM. 2014. A comparison of dual gradient-echo and spin-echo fMRI of the inferior temporal lobe. Human Brain Mapping. 35:4118–4128.

Harry BB, Umla-Runge K, Lawrence AD, Graham KS, Downing PE. 2016. Evidence for integrated visual face and body representations in the anterior temporal lobes. Journal of Cognitive Neuroscience.

Heleven E, van Overwalle F. 2018. The neural basis of representing others’ inner states. Current Opinion in Psychology.

Hennion S, Delbeuck X, Koelkebeck K, Brion M, Tyvaert L, Plomhause L, Derambure P, Lopes R, Szurhaj W. 2016. A functional magnetic resonance imaging investigation of theory of mind impairments in patients with temporal lobe epilepsy. Neuropsychologia. 93:271–279.

Hodges JR, Patterson K. 2007. Semantic dementia: a unique clinicopathological syndrome. The Lancet Neurology. 6:1004–1014.

Hoffman P, Binney RJ, Lambon Ralph MA. 2015. Differing contributions of inferior prefrontal and anterior temporal cortex to concrete and abstract conceptual knowledge. Cortex.

Hoffman P, Pobric G, Drakesmith M, Lambon Ralph MA. 2012. Posterior middle temporal gyrus is involved in verbal and non-verbal semantic cognition: Evidence from rTMS. Aphasiology. 26:1119–1130.

Humphreys GF, Jackson RL, Lambon Ralph MA. 2020. Overarching Principles and Dimensions of the Functional Organization in the Inferior Parietal Cortex. Cerebral Cortex. 30:5639–5653.

Irish M, Hodges JR, Piguet O. 2014. Right anterior temporal lobe dysfunction underlies theory of mind impairments in semantic dementia. Brain. 137:1241–1253.

Jacoby N, Bruneau E, Koster-Hale J, Saxe R. 2016. Localizing Pain Matrix and Theory of Mind networks with both verbal and non-verbal stimuli. NeuroImage. 126:39–48.

Jefferies E, Lambon Ralph MA. 2006. Semantic impairment in stroke aphasia versus semantic dementia: A case-series comparison. Brain. 129:2132–2147.

Kana RK, Keller T, Cherkassky VL, Minshew NJ, Just MA. 2009. Atypical frontal-posterior synchronization of Theory of Mind regions in autism during mental state attribution. Social Neuroscience. 4:135–152.

Klüver H, Bucy PC. 1937. “ Psychic blindness” and other symptoms following bilateral temporal lobectomy in Rhesus monkeys. American Journal of Physiology. 119:3552–353.

Kumfor F, Hazelton JL, de Winter F-L, de Langavant LC, van den Stock J. 2017. Clinical Studies of Social Neuroscience: A Lesion Model Approach. In: Neuroscience and Social Science. Cham: Springer International Publishing. p. 255–296.

Kumfor F, Honan C, McDonald S, Hazelton JL, Hodges JR, Piguet O. 2017. Assessing the “social brain” in dementia: Applying TASIT-S. Cortex. 93:166–177.

Kumfor F, Irish M, Hodges JR, Piguet O. 2013. Discrete Neural Correlates for the Recognition of Negative Emotions: Insights from Frontotemporal Dementia. PLoS ONE. 8.

Kumfor F, Landin-Romero R, Devenney E, Hutchings R, Grasso R, Hodges JR, Piguet O. 2016. On the right side? A longitudinal study of left-versus right-lateralized semantic dementia. Brain. 139:986–998.

Kumfor F, Piguet O. 2012. Disturbance of emotion processing in frontotemporal dementia: A synthesis of cognitive and neuroimaging findings. Neuropsychology Review.

Lambon Ralph MA, Mcclelland JL, Patterson K, Galton CJ, Hodges JR. 2001. No right to speak? The relationship between object naming and semantic impairment: Neuropsychological evidence and a computational model. Journal of Cognitive Neuroscience. 13:341–356.

Lieberman MD. 2007. Social Cognitive Neuroscience: A Review of Core Processes. Annual Review of Psychology. 58:259–289.

Lin N, Wang X, Xu Y, Wang X, Hua H, Zhao Y, Li X. 2018. Fine Subdivisions of the Semantic Network Supporting Social and Sensory–Motor Semantic Processing. Cerebral Cortex. 28:2699–2710.

Lin N, Yang X, Li J, Wang S, Hua H, Ma Y, Li X. 2018. Neural correlates of three cognitive processes involved in theory of mind and discourse comprehension. Cognitive, Affective and Behavioral Neuroscience. 18:273–283.

Luzzi S, Snowden JS, Neary D, Coccia M, Provinciali L, Lambon Ralph MA. 2007. Distinct patterns of olfactory impairment in Alzheimer’s disease, semantic dementia, frontotemporal dementia, and corticobasal degeneration. Neuropsychologia. 45:1823–1831.

Marinkovic K, Dhond RP, Dale AM, Glessner M, Carr V, Halgren E. 2003. Spatiotemporal Dynamics of Modality-Specific and Supramodal Word Processing The importance of the left anterior temporal cortex for. Neuron. 38:487–497.

Mellem MS, Jasmin KM, Peng C, Martin A. 2016. Sentence processing in anterior superior temporal cortex shows a social-emotional bias. Neuropsychologia. 89:217–224.

Mion M, Patterson K, Acosta-Cabronero J, Pengas G, Izquierdo-Garcia D, Hong YT, Fryer TD, Williams GB, Hodges JR, Nestor PJ. 2010. What the left and right anterior fusiform gyri tell us about semantic memory. Brain. 133:3256–3268.

Molenberghs P, Johnson H, Henry JD, Mattingley JB. 2016. Understanding the minds of others: A neuroimaging meta-analysis. Neuroscience and Biobehavioral Reviews. 65:276–291.

Moll J, Zahn R, de Oliveira-Souza R, Krueger F, Grafman J. 2005. Opinion: The neural basis of human moral cognition. Nature Reviews Neuroscience.

Mummery CJ, Patterson K, Price CJ, Ashburner J, Frackowiak RSJ, Hodges JR. 2000. A voxel-based morphometry study of semantic dementia: Relationship between temporal lobe atrophy and semantic memory. Annals of Neurology. 47:36–45.

Munafò MR, Nosek BA, Bishop DVM, Button KS, Chambers CD, Percie Du Sert N, Simonsohn U, Wagenmakers EJ, Ware JJ, Ioannidis JPA. 2017. A manifesto for reproducible science. Nature Human Behaviour.

Nestor PJ, Fryer TD, Hodges JR. 2006. Declarative memory impairments in Alzheimer’s disease and semantic dementia. NeuroImage. 30:1010–1020.

Nichols T, Brett M, Andersson J, Wager T, Poline JB. 2005a. Valid conjunction inference with the minimum statistic. NeuroImage. 25:653–660.

Nichols T, Brett M, Andersson J, Wager T, Poline JB. 2005b. Valid conjunction inference with the minimum statistic. NeuroImage. 25:653–660.

Oldfield RC. 1971. The assessment and analysis of handedness: The Edinburgh inventory. Neuropsychologia. 9:97–113.

Olson IR, McCoy D, Klobusicky E, Ross LA. 2013. Social cognition and the anterior temporal lobes: A review and theoretical framework. Social Cognitive and Affective Neuroscience.

Olson IR, Plotzker A, Ezzyat Y. 2007. The Enigmatic temporal pole: A review of findings on social and emotional processing. Brain.

Papinutto N, Galantucci S, Mandelli ML, Gesierich B, Jovicich J, Caverzasi E, Henry RG, Seeley WW, Miller BL, Shapiro KA, Gorno-Tempini ML. 2016. Structural connectivity of the human anterior temporal lobe: A diffusion magnetic resonance imaging study. Human Brain Mapping. 37:2210–2222.

Pascual B, Masdeu JC, Hollenbeck M, Makris N, Insausti R, Ding SL, Dickerson BC. 2015. Large-scale brain networks of the human left temporal pole: A functional connectivity MRI study. Cerebral Cortex. 25:680–702.

Patterson K, Nestor PJ, Rogers TT. 2007. Where do you know what you know? The representation of semantic knowledge in the human brain. Nature Reviews Neuroscience.

Pelli DG. 1997. The VideoToolbox software for visual psychophysics: transforming numbers into movies. Spatial Vision. 10:437–442.

Persichetti AS, Denning JM, Gotts SJ, Martin A. 2021. A Data-Driven Functional Mapping of the Anterior Temporal Lobes. The Journal of Neuroscience. 41:6038–6049.

Piwnica Worms KE, Omar R, Hailstone JC, Warren JD. 2010. Flavour processing in semantic dementia. Cortex. 46:761–768.

Plaut DC. 2002. Graded modality-specific specialisation in semantics: A computational account of optic aphasia, Cognitive Neuropsychology.

Pobric G, Jefferies E, Ralph AL. 2008. Anterior temporal lobes and non-verbal semantic processing: Brain Stimulation. 1:307.

Pobric G, Ralph MAL, Zahn R. 2016. Hemispheric specialization within the superior anterior temporal cortex for social and nonsocial concepts. Journal of Cognitive Neuroscience. 28:351–360.

Price CJ, Friston KJ. 1997. Cognitive conjunction: A new approach to brain activation experiments. NeuroImage. 5:261–270.

Ralph MAL, Jefferies E, Patterson K, Rogers TT. 2017. The neural and computational bases of semantic cognition. Nature Reviews Neuroscience. 18:42–55.

Ramot M, Walsh C, Martin A. 2019. Multifaceted integration: Memory for faces is subserved by widespread connections between visual, memory, auditory, and social networks. Journal of Neuroscience. 39:4976–4985.

Ramsey R, Ward R. 2020. Putting the Nonsocial Into Social Neuroscience: A Role for Domain-General Priority Maps During Social Interactions. Perspectives on Psychological Science. 15:1076–1094.

Rankin KP. 2020. Brain Networks Supporting Social Cognition in Dementia. Current Behavioral Neuroscience Reports. 203–211.

Rankin KP. 2021. Measuring Behavior and Social Cognition in FTLD. In: Frontotemporal Dementias. Springer. p. 51–65.

Rice GE, Caswell H, Moore P, Hoffman P, Lambon Ralph MA. 2017. The effects of left versus right anterior temporal lobe resection on semantic processing of words, objects, and faces. Journal of the Neurological Sciences. 381:684.

Rice GE, Hoffman P, Binney RJ, Lambon Ralph MA. 2018. Concrete versus abstract forms of social concept: An fMRI comparison of knowledge about people versus social terms. Philosophical Transactions of the Royal Society B: Biological Sciences. 373:20170136.

Rice GE, Hoffman P, Lambon Ralph MA. 2015. Graded specialization within and between the anterior temporal lobes. Annals of the New York Academy of Sciences. 1359:84–97.

Rice GE, Ralph MAL, Hoffman P. 2015. The roles of left versus right anterior temporal lobes in conceptual knowledge: An ALE meta-analysis of 97 functional neuroimaging studies. Cerebral Cortex. 25:4374–4391.

Ross LA, Olson IR. 2010. Social cognition and the anterior temporal lobes. NeuroImage. 49:3452–3462.

Saxe R. 2006. Why and how to study Theory of Mind with fMRI. Brain Research. 1079:57–65.

Saxe R, Kanwisher N. 2003a. People thinking about thinking people. The role of the temporo-parietal junction in &quot;theory of mind&quot;. NeuroImage. 19:1835–1842.

Saxe R, Kanwisher N. 2003b. People thinking about thinking peopleThe role of the temporo-parietal junction in “theory of mind.” NeuroImage. 19:1835–1842.

Saxe R, Powell LJ. 2006. It’s the Thought That Counts: Specific Brain Regions for One Component of Theory of Mind, Source: Psychological Science.

Saxe R, Wexler A. 2005. Making sense of another mind: the role of the right temporo-parietal junction. Neuropsychologia. 43:1391–1399.

Scholz J, Triantafyllou C, Whitfield-Gabrieli S, Brown EN, Saxe R. 2009. Distinct regions of right temporo-parietal junction are selective for theory of mind and exogenous attention. PLoS ONE. 4.

Schurz M, Radua J, Aichhorn M, Richlan F, Perner J. 2014. Fractionating theory of mind: A meta-analysis of functional brain imaging studies. Neuroscience & Biobehavioral Reviews. 42:9–34.

Seghier ML, Fagan E, Price CJ. 2010. Functional Subdivisions in the Left Angular Gyrus Where the Semantic System Meets and Diverges from the Default Network. Journal of Neuroscience. 30:16809–16817.

Shimotake A, Matsumoto R, Ueno T, Kunieda T, Saito S, Hoffman P, Kikuchi T, Fukuyama H, Miyamoto S, Takahashi R, Ikeda A, Lambon Ralph MA. 2015. Direct exploration of the role of the ventral anterior temporal lobe in semantic memory: Cortical stimulation and local field potential evidence from subdural grid electrodes. Cerebral Cortex. 25:3802–3817.

Simmons WK, Martin A. 2009. The anterior temporal lobes and the functional architecture of semantic memory. Journal of the International Neuropsychological Society. 15:645–649.

Simmons WK, Reddish M, Bellgowan PSFF, Martin A. 2010. The selectivity and functional connectivity of the anterior temporal lobes. Cerebral Cortex. 20:813–825.

Snowden JS, Harris JM, Thompson JC, Kobylecki C, Jones M, Richardson AM, Neary D. 2018. Semantic dementia and the left and right temporal lobes. Cortex. 107:188–203.

Spitsyna G, Warren JE, Scott SK, Turkheimer FE, Wise RJS. 2006. Converging language streams in the human temporal lobe. Journal of Neuroscience. 26:7328–7336.

Spunt RP, Adolphs R. 2017. A new look at domain specificity: Insights from social neuroscience. Nature Reviews Neuroscience.

Synn A, Mothakunnel A, Kumfor F, Chen Y, Piguet O, Hodges JR, Irish M. 2018. Mental states in moving shapes: distinct cortical and subcortical contributions to theory of mind impairments in dementia. Journal of Alzheimer’s Disease. 61:521–535.

Terzian H, Dalle Ore G. 1955. Syndrome of Klüver and Bucy, reproduced in man by bilateral removal of the temporal lobes. Neurology.

van Hoeck N, Begtas E, Steen J, Kestemont J, Vandekerckhove M, van Overwalle F. 2014. False belief and counterfactual reasoning in a social environment. NeuroImage. 90:315–325.

van Overwalle F. 2009. Social cognition and the brain: A meta-analysis. Human Brain Mapping. 30:829–858.

Vigliocco G, Kousta S-TT, della Rosa PA, Vinson DP, Tettamanti M, Devlin JT, Cappa SF. 2014. The neural representation of abstract words: The role of emotion. Cerebral Cortex. 24:1767–1777.

Visser M, Embleton K v., Jefferies E, Parker GJ, Ralph MAL. 2010. The inferior, anterior temporal lobes and semantic memory clarified: Novel evidence from distortion-corrected fMRI. Neuropsychologia. 48:1689–1696.

Visser M, Jefferies E, Embleton K v., Lambon Ralph MA. 2012. Both the Middle Temporal Gyrus and the Ventral Anterior Temporal Area Are Crucial for Multimodal Semantic Processing: Distortion-corrected fMRI Evidence for a Double Gradient of Information Convergence in the Temporal Lobes. Journal of Cognitive Neuroscience. 24:1766–1778.

Visser M, Jefferies E, Lambon Ralph MA, Ralph L, Visser M, Jefferies E, Lambon Ralph MA. 2010. Semantic Processing in the Anterior Temporal Lobes : A Meta-analysis of the Functional. Journal of Cognitive Neuroscience. 22:1083–1094.

Visser M, Lambon Ralph MA. 2011. Differential Contributions of Bilateral Ventral Anterior Temporal Lobe and Left Anterior Superior Temporal Gyrus to Semantic Processes. Journal of Cognitive Neuroscience. 23:3121–3131.

Walbrin J, Downing P, Koldewyn K. 2018. Neural responses to visually observed social interactions. Neuropsychologia. 112:31–39.

Wang X, Wang B, Bi Y. 2019. Close yet independent: Dissociation of social from valence and abstract semantic dimensions in the left anterior temporal lobe. Human Brain Mapping. 40:4759–4776.

Wong C, Gallate J. 2012. The function of the anterior temporal lobe: A review of the empirical evidence. Brain Research. 1449:94–116.

Young L, Dodell-Feder D, Saxe R. 2010. What gets the attention of the temporo-parietal junction? An fMRI investigation of attention and theory of mind. Neuropsychologia. 48:2658–2664.

Zahn R, Moll J, Iyengar V, Huey ED, Tierney M, Krueger F, Grafman J. 2009. Social conceptual impairments in frontotemporal lobar degeneration with right anterior temporal hypometabolism. Brain. 132:604–616.

Zahn R, Moll J, Krueger F, Huey ED, Garrido G, Grafman J. 2007. Social concepts are represented in the superior anterior temporal cortex. Proceedings of the National Academy of Sciences. 104:6430–6435.

