## Supplementary Information for "The Role of the Ventrolateral Anterior Temporal Lobes in Social Cognition"

for

#### Supplementary Figures

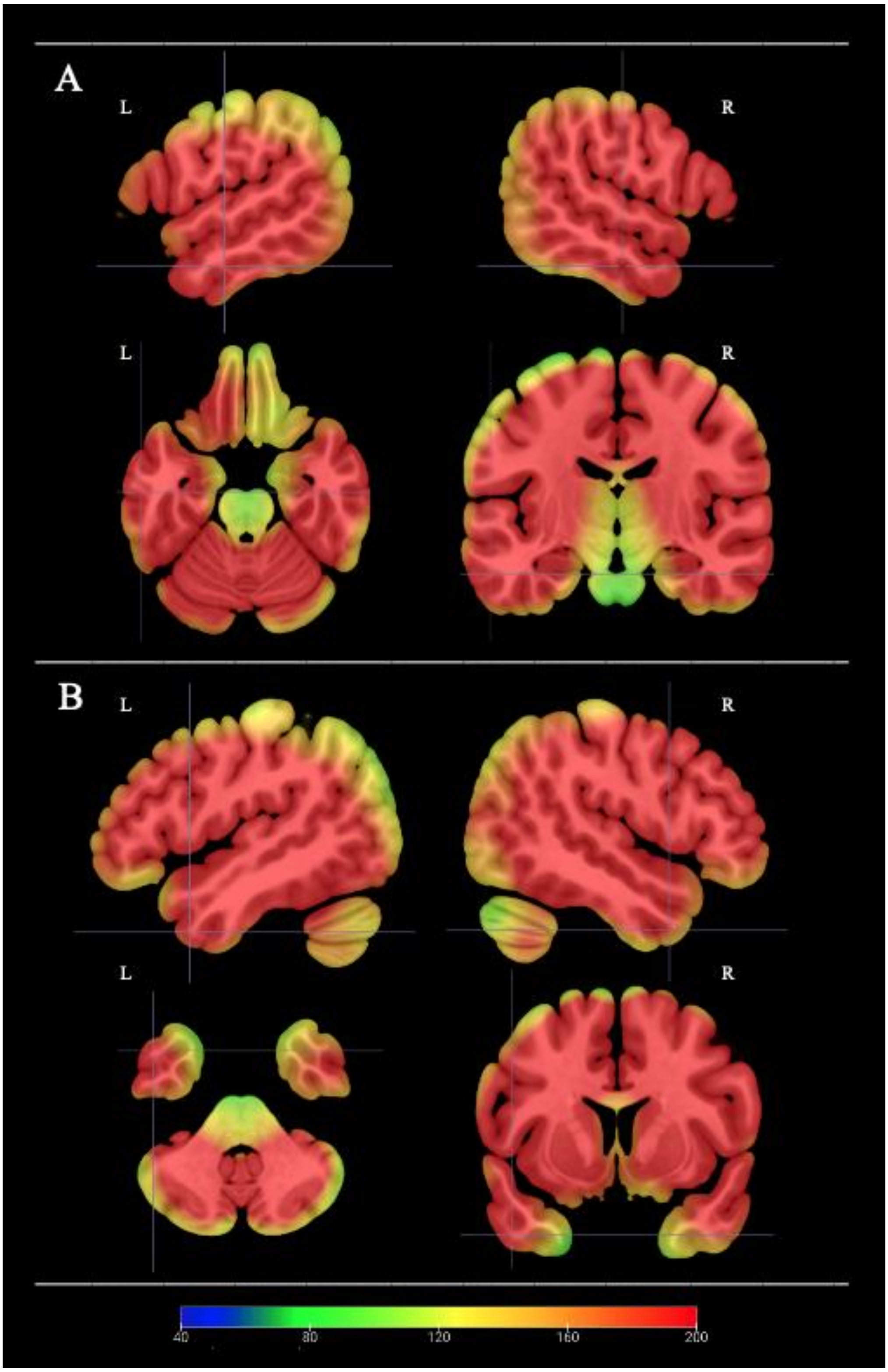

***Supplementary Figure M1.*** *Temporal signal-to-noise ratio (tSNR) maps showing dual echo EPI image quality over the anterior temporal lobes. The colour gradient indicates the tSNR of the pre-processed EPI time course data overlaid on a MNI template. tSNR was calculated for each functional run in each participant by dividing the mean intensity in each voxel by its standard deviation. The resulting tSNR maps displayed here were averaged across runs and participants. The colour map is thresholded at tSNR of 40, with all areas in red indicating a tSNR of at least 200. Cross-sections were chosen to display (Panel A) a ventrolateral ATL region implicated in general semantic processing (Visser, Jefferies, Embleton & Lambon Ralph, 2012 [-57,* ***-15****, -24]), and (B) a polar ATL region implicated in the processing of social concepts (Binney, Hoffman & Lambon Ralph, 2016 [-48,* ***9****, -39]).*

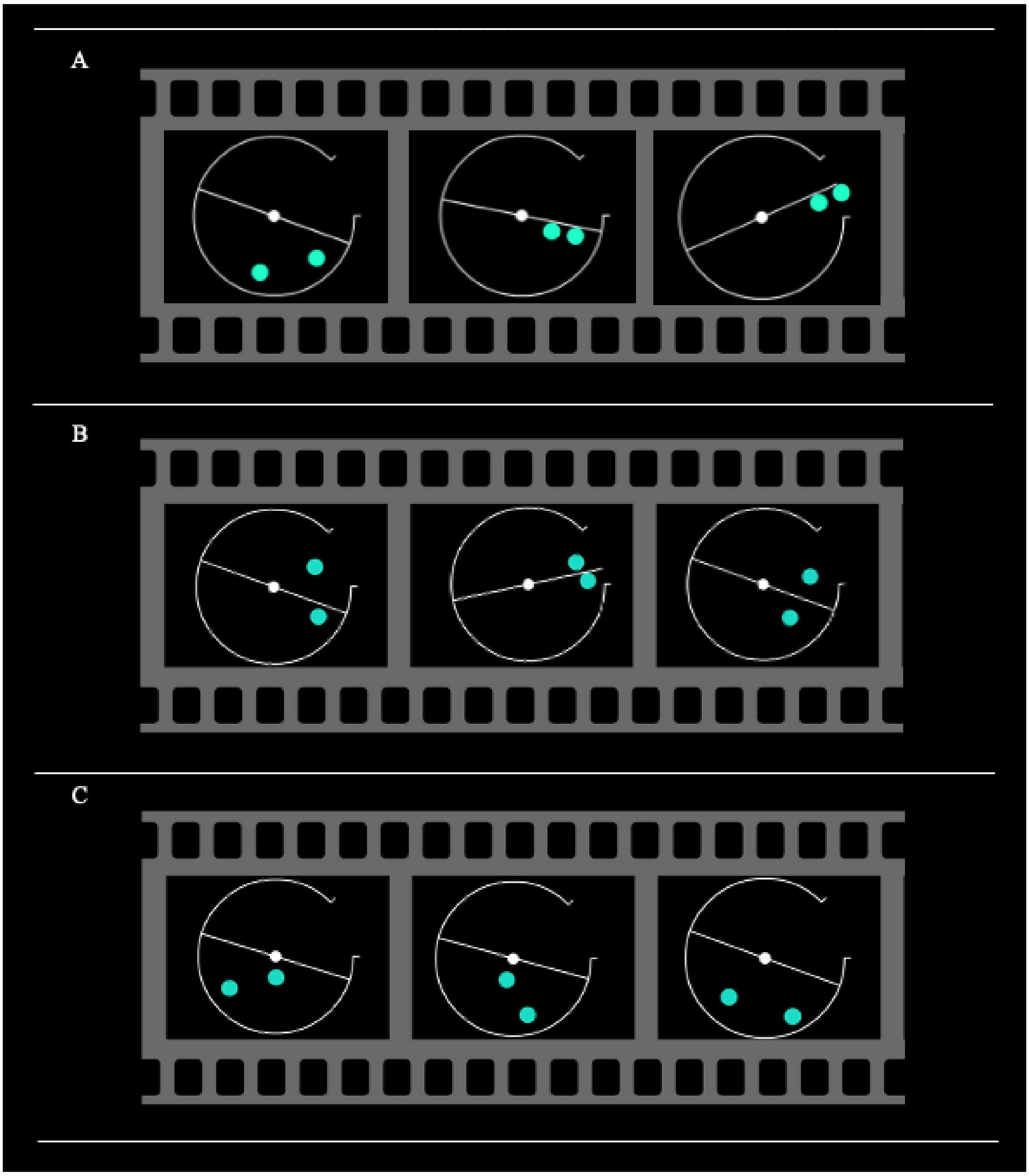

***Supplementary Figure M2.*** *Schematic examples of the main experimental interaction judgement theory of mind task stimuli. Panel A) displays a friendly (cooperative) interaction. Panel B) displays an unfriendly (competitive) interaction. Panel C) displays a control stimulus in which the shapes move randomly at same or different speeds. The .avi files of the actual stimuli examples can be accessed via OSF ([https://osf.io/v2gt5/](https://osf.io/v2gt5/)).*

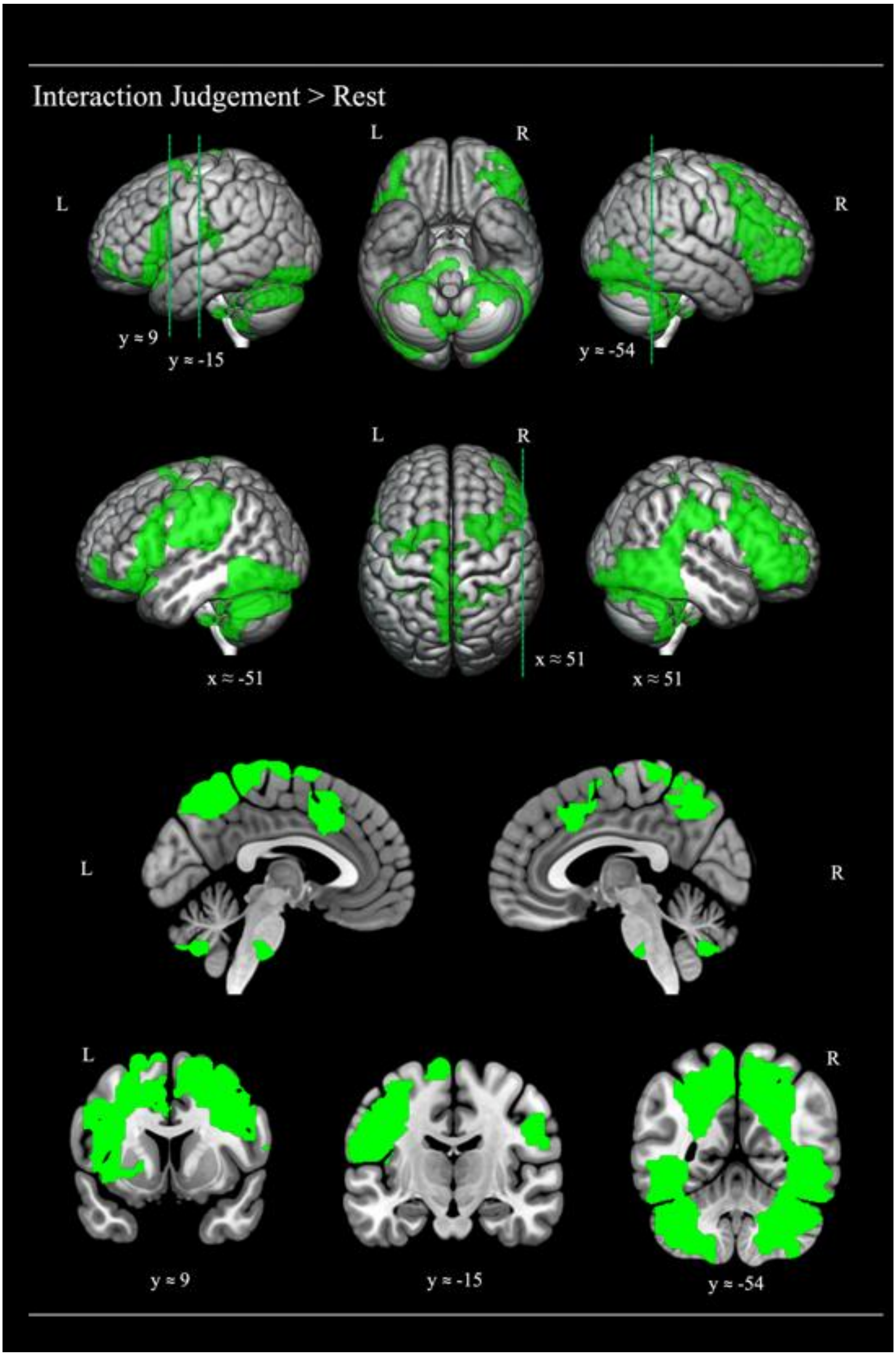

***Supplementary Figure 1.*** *Cortical regions activated by the social interaction judgement condition, relative to rest. The statistical map was thresholded with an uncorrected voxel height threshold of p < .001 and a family wise error corrected minimum cluster extent threshold (k= 17513) at p < .05. Cross-sections were chosen to display the location of activation found in key studies investigating ToM processing (Saxe & Kanwisher, 2003; right TPJ [**51**, -**54**, 27]), semantic processing of social concepts (Binney, Hoffman & Lambon Ralph, 2016; left TP [-48, **9**, -39]) and general semantic processing (Visser, Jefferies, Embleton & Lambon Ralph, 2012; left inferior ATL [-57, -**15**, -24]).*

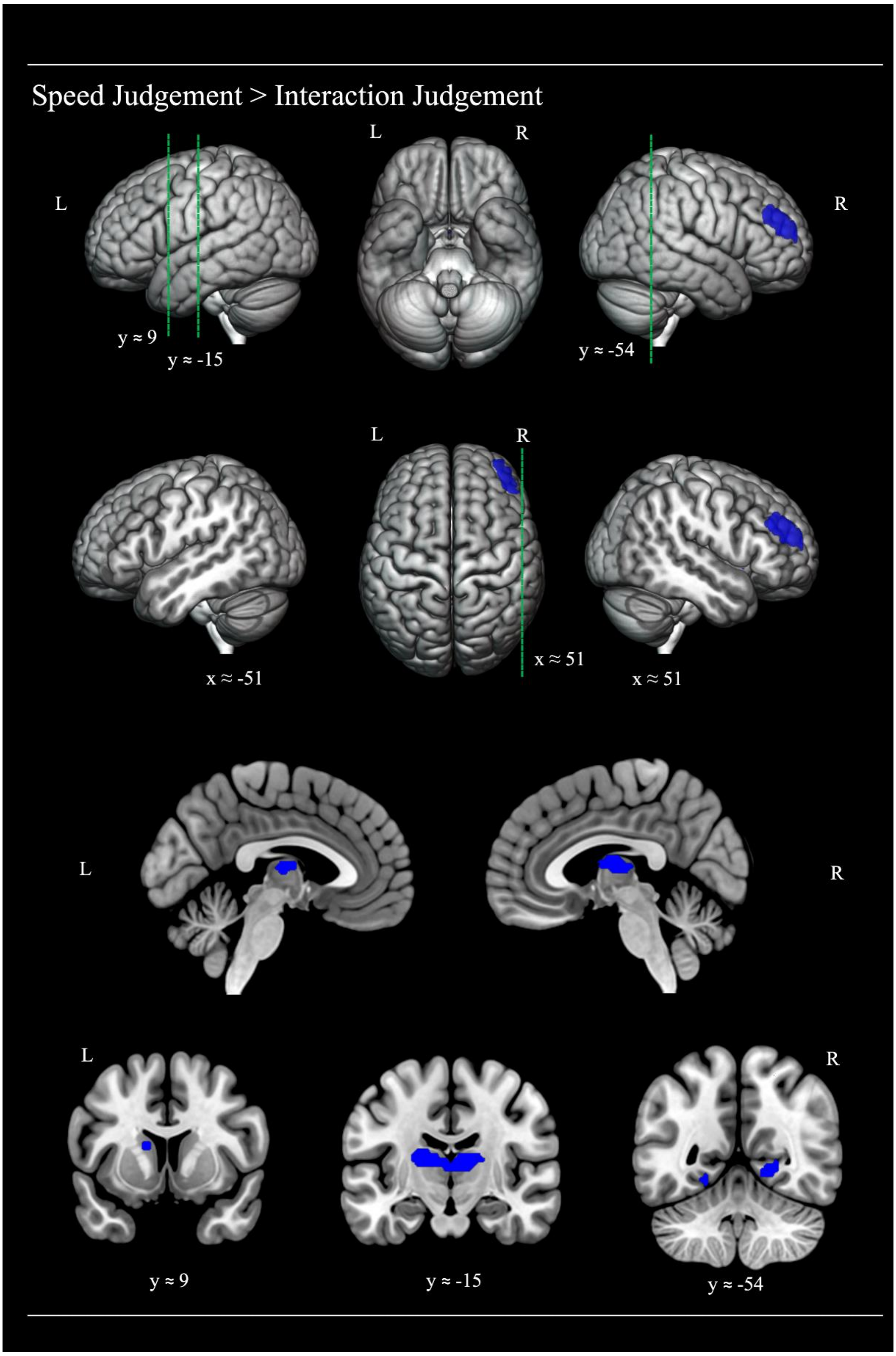

***Supplementary Figure 2.*** *Cortical regions activated by the speed judgement control condition, relative to the social interaction judgement condition. The statistical map was thresholded with an uncorrected voxel height threshold of p < .001 and a family wise error corrected minimum cluster extent threshold (k= 125) at p < .05. Cross-sections were chosen to display the location of activation found in key studies investigating ToM processing (Saxe & Kanwisher, 2003; right TPJ [**51**, **-54**, 27]), semantic processing of social concepts (Binney, Hoffman & Lambon Ralph, 2016; left TP [-48, **9**, -39]) and general semantic processing (Visser, Jefferies, Embleton & Lambon Ralph, 2012; left inferior ATL [-57, -**15**, -24]).*

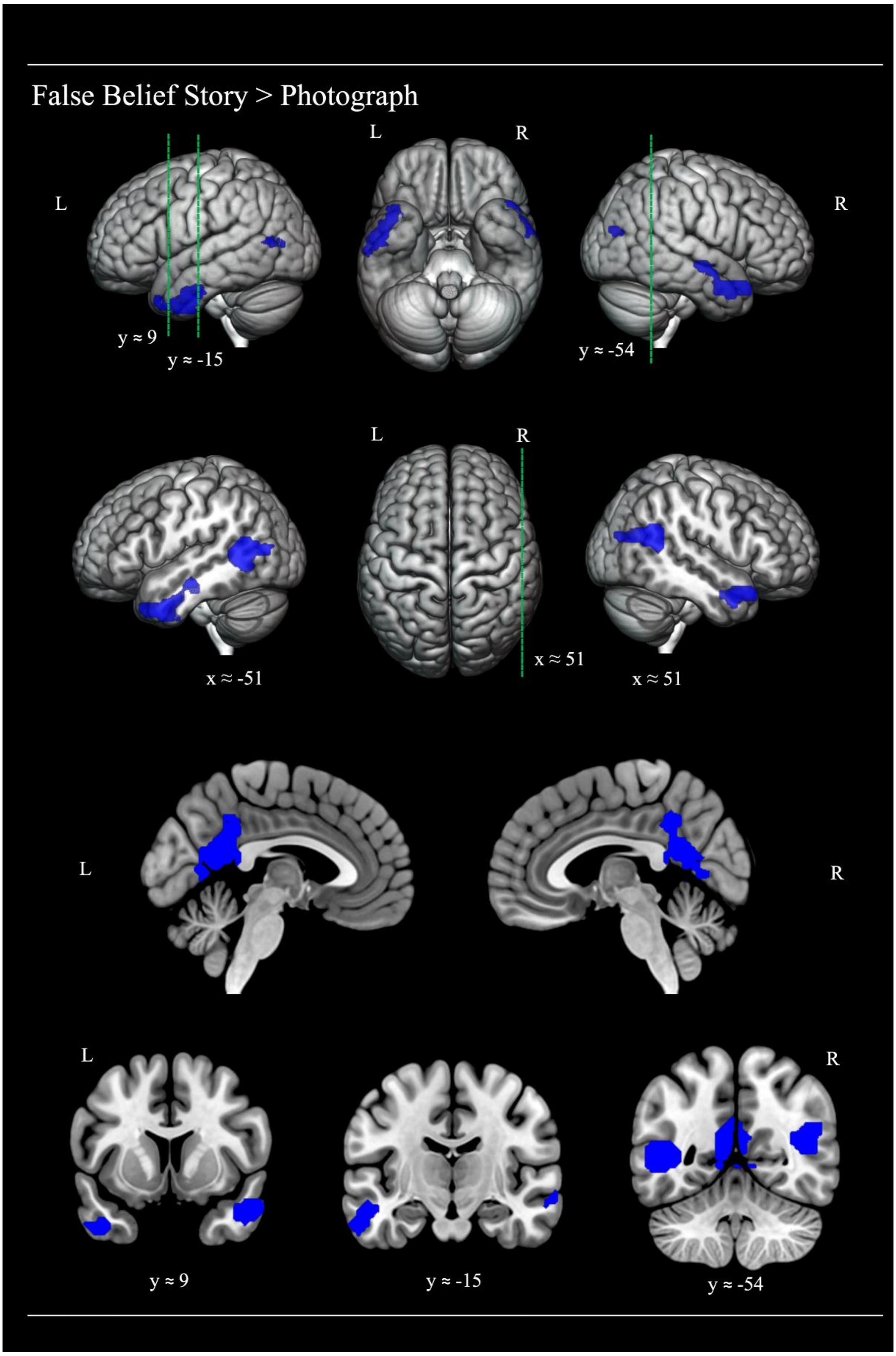

***Supplementary Figure 3.*** *Cortical regions activated by the false belief condition, relative to the false photograph control condition. The statistical map was thresholded with an uncorrected voxel height threshold of p < .001 and a family wise error corrected minimum cluster extent threshold (k= 247) at p < .05. Cross-sections were chosen to display the location of activation found in key studies investigating ToM processing (Saxe & Kanwisher, 2003; right TPJ [**51**, **-54**, 27]), semantic processing of social concepts (Binney, Hoffman & Lambon Ralph, 2016; left TP [-48, **9**, -39]) and general semantic processing (Visser, Jefferies, Embleton & Lambon Ralph, 2012; left inferior ATL [-57, **-15**, -24]).*

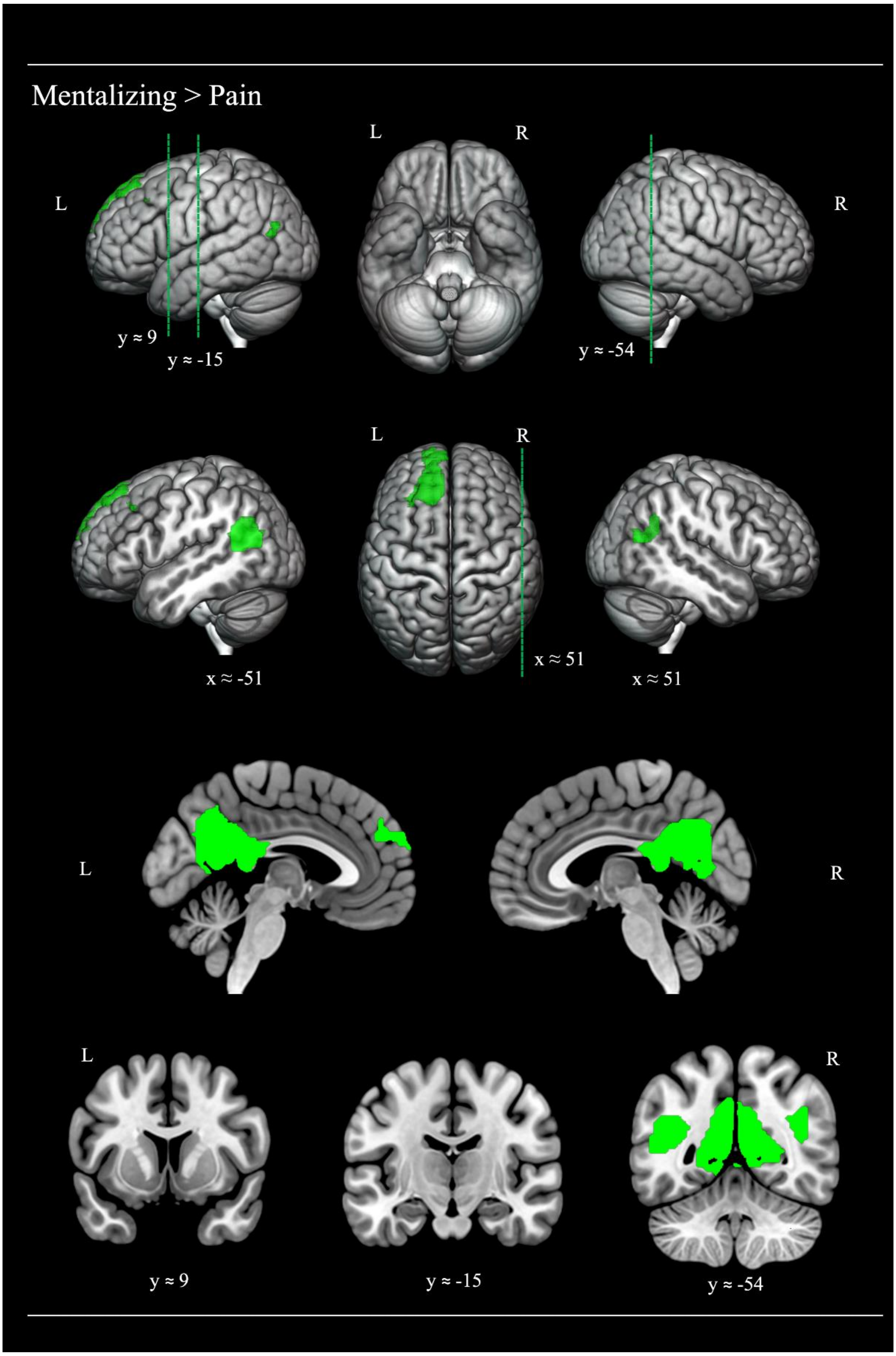

***Supplementary Figure 4.*** *Cortical regions revealed by the mentalising > pain contrast performed upon the free-viewing movie ToM paradigm. The statistical map was thresholded with an uncorrected voxel height threshold of p < .001 and a family wise error corrected minimum cluster extent threshold (k= 163) at p < .05. Cross-sections were chosen to display the location of activation found in key studies investigating ToM processing (Saxe & Kanwisher, 2003; right TPJ [**51**, -**54**, 27]), semantic processing of social concepts (Binney, Hoffman & Lambon Ralph, 2016; left TP [-48, **9**, -39]) and general semantic processing (Visser, Jefferies, Embleton & Lambon Ralph, 2012; left inferior ATL [-57, -**15**, -24]).*

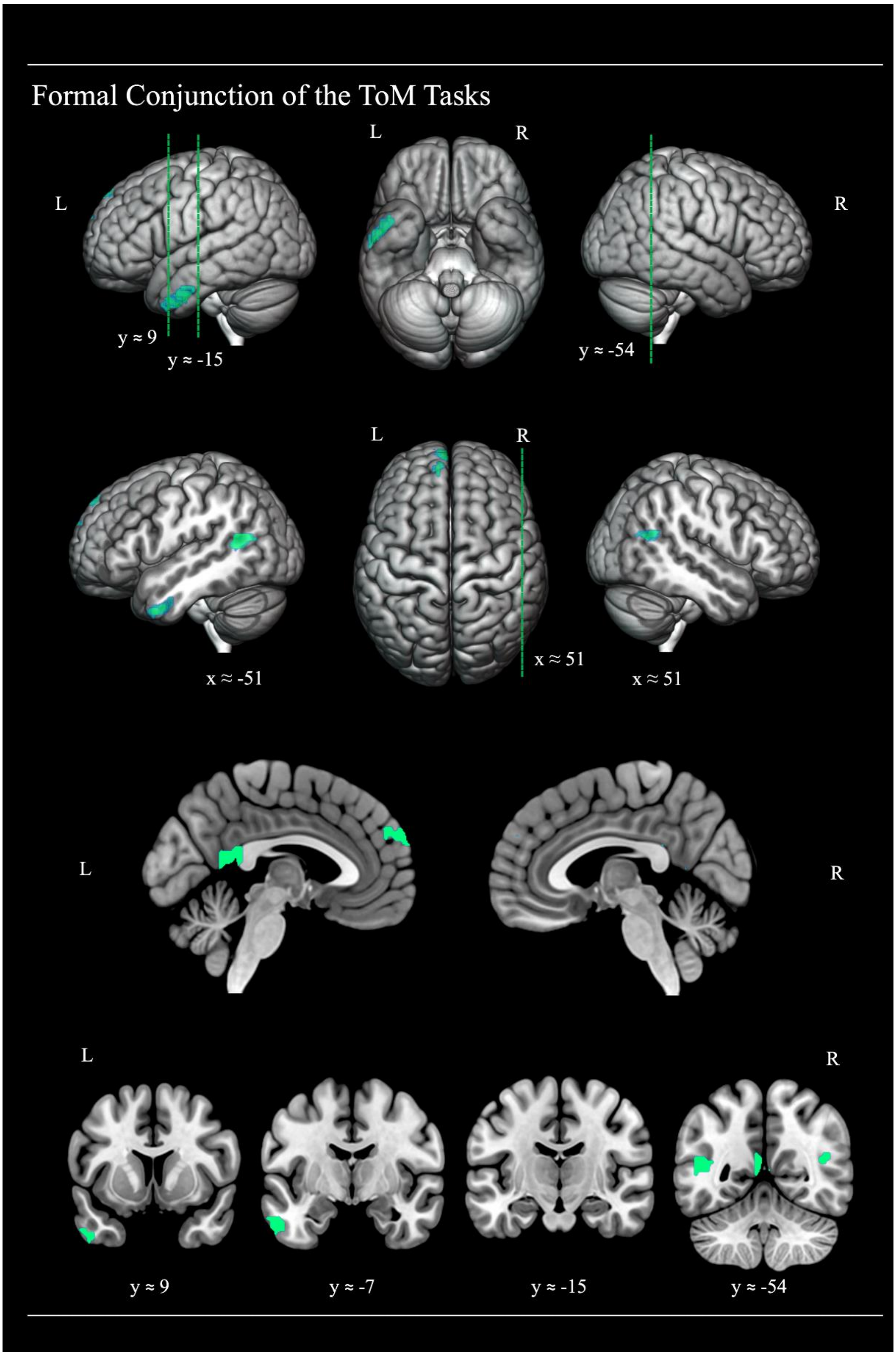

***Supplementary Figure 5.*** *Common activation of cortical regions by the interaction judgement > speed judgement contrast of the main experimental IJ ToM task and the false belief story > photograph contrast of the FB ToM localiser and the mentalising > pain contrast of the free viewing movie ToM localiser. The statistical maps were thresholded with an uncorrected voxel height threshold of p < .001. Cross-sections were chosen to display the location of activation found in key studies investigating ToM processing (Saxe & Kanwisher, 2003; right TPJ [**51**, **-54**, 27]), semantic processing of social concepts (Binney, Hoffman & Lambon Ralph, 2016; left TP [-48, **9**, -39]) and general semantic processing (Visser, Jefferies, Embleton & Lambon Ralph, 2012; left inferior ATL [-57, **-15**, -24]), as well one further key area of 3-way overlap (y = -7).*

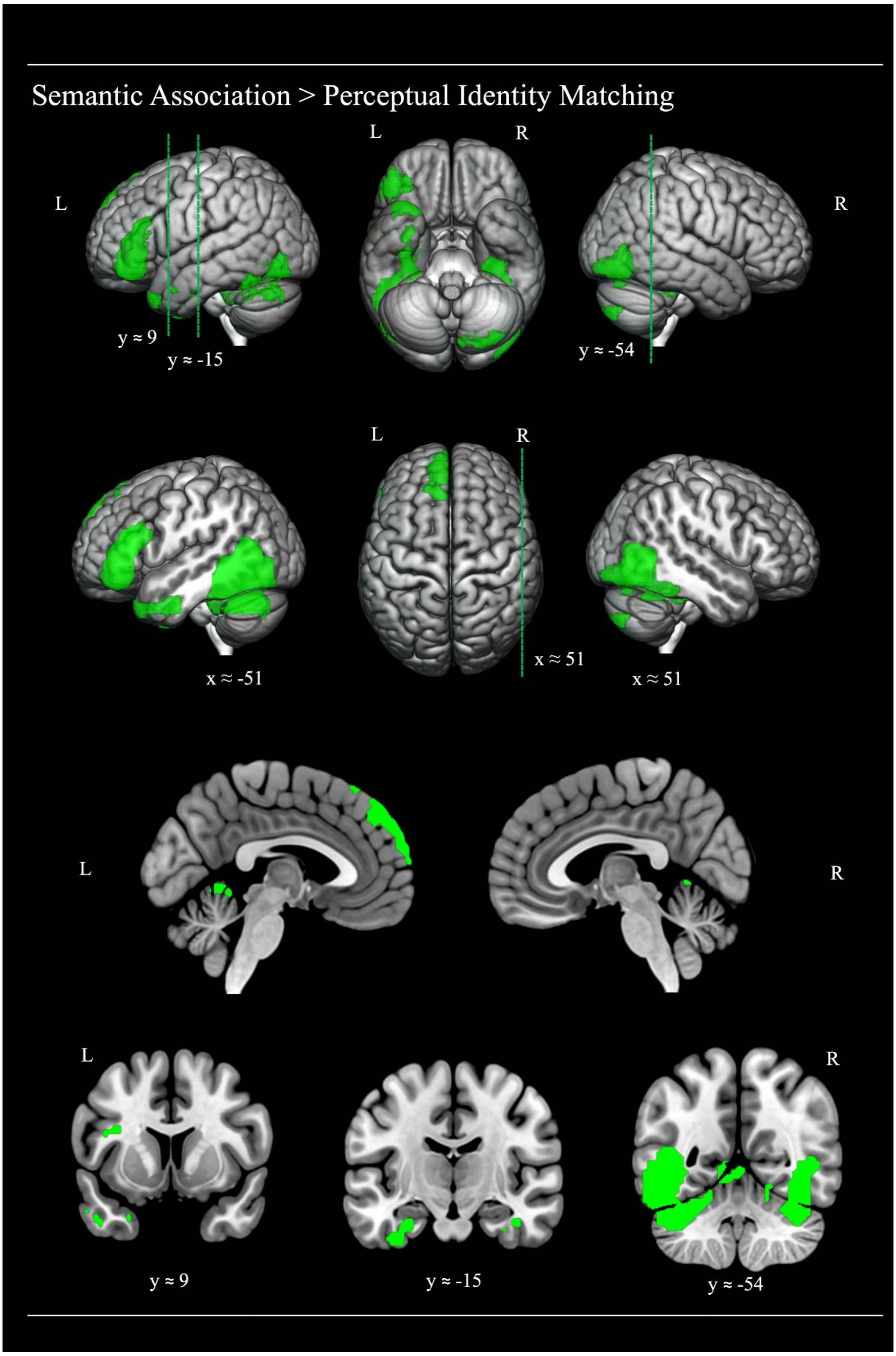

***Supplementary Figure 6.*** *Cortical regions activated by the semantic association task relative to the perceptual matching control condition. The statistical map was thresholded with an uncorrected voxel height threshold of p < .001 and a family wise error corrected minimum cluster extent threshold (k= 288) at p < .05. Cross-sections were chosen to display the location of activation found in key studies investigating ToM processing (Saxe & Kanwisher, 2003; right TPJ [**51**, -**54**, 27]), semantic processing of social concepts (Binney, Hoffman & Lambon Ralph, 2016; left TP [-48, **9**, -39]) and general semantic processing (Visser, Jefferies, Embleton & Lambon Ralph, 2012; left inferior ATL [-57, -**15**, -24]).*

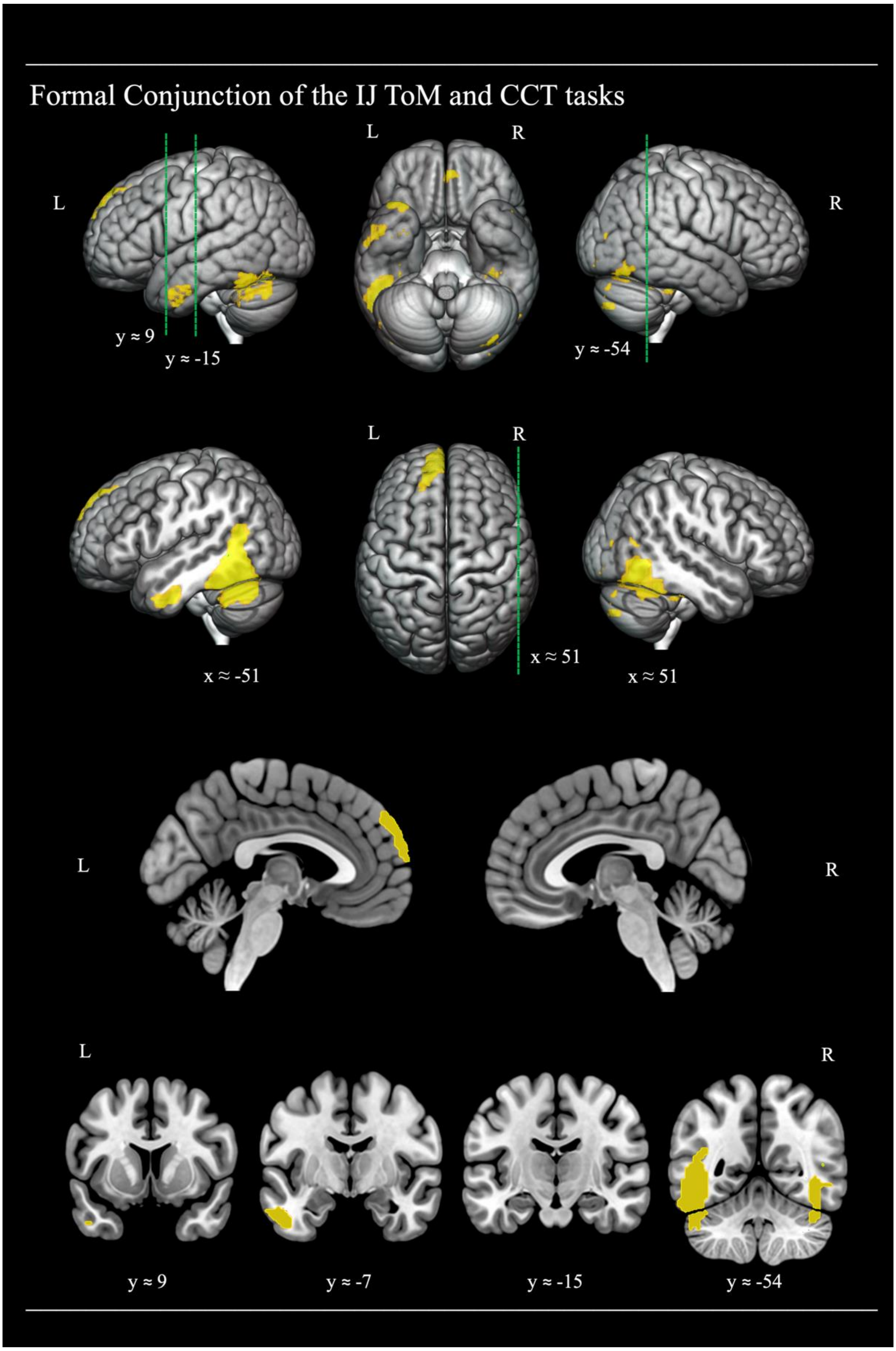

***Supplementary Figure 7.*** *Common activation of cortical regions by the interaction judgement > speed judgement contrast of the main experimental IJ ToM task and the semantic judgement > perceptual judgement contrast of the CCT semantic task. The statistical maps were thresholded with an uncorrected voxel height threshold of p < .001. Cross-sections were chosen to display the location of activation found in key studies investigating ToM processing (Saxe & Kanwisher, 2003; right TPJ [***51***,* ***-54****, 27]), semantic processing of social concepts (Binney, Hoffman & Lambon Ralph, 2016; left TP [-48,* ***9****, -39]) and general semantic processing (Visser, Jefferies, Embleton & Lambon Ralph, 2012; left inferior ATL [-57,* ***-15****, -24]), as well one further key area of 3-way overlap (y = -7)*

### Supplementary Tables

***Supplementary Table M1*** Family-wise error corrected cluster extent thresholds (FWEc), smoothness of data, search volume and number of RESELS for each univariate contrast

| Contrast | FWEc | Smoothness | | | Search Volume (voxels) | RESELS |
|---|---|---|---|---|---|---|
| **Theory of Mind Task** | | | | | | |
| Interaction > Speed | 152 | 16.5 | 16.5 | 14.8 | 67502 | 3863 |
| Speed > Interaction | 125 | 16.5 | 16.5 | 14.8 | | 3863 |
| Interaction > Rest | 17513 | 15.8 | 15.3 | 14.7 | | 4430 |
| **Camel & Cactus Task** | | | | | | |
| Semantic > Perceptual | 288 | 16 | 15.8 | 14.3 | 67502 | 4303 |
| **False Belief localiser** | | | | | | |
| False Belief > False Fact | 247 | 18 | 18 | 14.6 | 67502 | 3324 |
| **Pixar Movie localiser** | | | | | | |
| Mentalizing > Pain | 163 | 17.3 | 16.6 | 14.9 | 67323 | 3616 |

***Supplementary Table R1*** Significant activation clusters in the interaction judgement > rest contrast (p < .05, FWE corrected with k= 17513 extent threshold and a cluster defining threshold of p< .001, uncorrected).

| **Cluster Name and Location of Maxima** | **Cluster Extent (voxels)** | **Peak (Z)** | **MNI Coordinates (mm)** | | |
|---|---|---|---|---|---|
| | | | **x** | **y** | **z** |
| L cerebellum | **17513** | 6.74 | -39 | -57 | -30 |
| L postcentral gyrus | | 6.18 | -48 | -21 | 30 |
| R occipito-temporal gyrus | | 6.13 | 45 | -63 | -12 |

*The table shows up to three local maxima per cluster more than 8.0 mm apart. L= left; R= right.*

***Supplementary Table R2*** Significant activation clusters in the speed judgement > interaction judgement contrast (p < .05, FWE corrected with k= 125 extent threshold and a cluster defining threshold of p< .001, uncorrected).

| **Cluster Name and Location of Maxima** | **Cluster Extent (voxels)** | **Peak (Z)** | **MNI Coordinates (mm)** | | |
|---|---|---|---|---|---|
| | | | **x** | **y** | **z** |
| R middle SFG | **247** | 4.96 | 36 | 42 | 24 |
| R middle SFG | | 4.04 | 36 | 60 | 18 |
| R hippocampus | **478** | 4.57 | 27 | -42 | 0 |
| R thalamus | | 4.26 | 6 | -9 | 12 |
| R lingual gyrus | | 4.26 | 21 | -57 | -3 |
| L lingual gyrus | **127** | 4.19 | -27 | -42 | -3 |
| L hippocampus | | 4.07 | -30 | -33 | -9 |
| L cerebellum | | 3.76 | -21 | -57 | -12 |
| R insula | **125** | 3.82 | 33 | 27 | 0 |
| R IFG | | 3.60 | 33 | 36 | 3 |

*The table shows up to three local maxima per cluster more than 8.0 mm apart. L= left; R= right. SFG = superior frontal gyrus; IFG = inferior frontal gyrus*

***Supplementary Table R3*** Planned comparisons across a priori ROIs for the social interaction > speed judgement contrast.

| Hemisphere | ROI pair | Mean (SD) | T-statistic | Sig. (2 tailed) | Effect Size (Cohen's d) |
|---|---|---|---|---|---|
| Left | vATL<br>TP | 0.22 (0.21)<br>0.18 (0.24) | 0.72 | .48 | 0.15 |
| | vATL<br>TPJ | 0.22 (0.21)<br>0.20 (0.24) | 0.33 | .74 | 0.07 |
| | TP<br>TPJ | 0.18 (0.24)<br>0.20 (0.24) | -0.19 | .85 | -0.04 |
| Right | vATL<br>TP | 0.11 (0.16)<br>0.23 (0.28) | -1.87 | .07 | -0.38 |
| | vATL<br>TPJ | 0.11 (0.16)<br>0.17 (0.19) | -1.2 | .24 | -0.24 |
| | TP<br>TPJ | 0.23 (0.28)<br>0.17 (0.19) | 0.9 | .38 | 0.18 |
| Left vs. Right | vATL (left)<br>vATL (right) | 0.22 (0.21)<br>0.11 (0.16) | 2.45 | .02 | 0.5 |
| | TP (left)<br>TP (right) | 0.18 (0.24)<br>0.23 (0.28) | -1.05 | .31 | -0.21 |
| | TPJ (left)<br>TPJ (right) | 0.20 (0.24)<br>0.17 (0.19) | 0.51 | .62 | 0.1 |

***Supplementary Table R4*** Significant activation clusters in the false belief > false photograph contrast ($p < .05$, FWE corrected with k= 247 extent threshold and a cluster defining threshold of $p < .001$, uncorrected).

| **Cluster Name and Location of Maxima** | **Cluster Extent (voxels)** | **Peak (Z)** | **MNI Coordinates (mm)** | | |
|---|---|---|---|---|---|
| | | | **x** | **y** | **z** |
| **L Posterior-Medial** | **492** | | | | |
| inferior precuneus | | 5.29 | -3 | -51 | 18 |
| lingual gyrus | | 3.9 | -9 | -57 | 3 |
| precuneus | | 3.89 | -3 | -48 | 36 |
| **R ATL** | **262** | | | | |
| superior TP | | 4.68 | 51 | 15 | -24 |
| anterior MTG | | 3.84 | 57 | 0 | -27 |
| superior TP | | 3.62 | 66 | -9 | -9 |
| **L Temporo-Occipital** | **367** | | | | |
| posterior MTG | | 4.42 | -48 | -57 | 9 |
| middle OG | | 3.52 | -45 | -75 | 9 |
| fusiform | | 3.32 | -39 | -60 | -12 |
| **L ATL** | **353** | | | | |
| anterior ITG | | 4.39 | -51 | 0 | -39 |
| anterior ITG | | 4.21 | -60 | -9 | -30 |
| anterior MTG / TP | | 3.83 | -42 | 9 | -36 |
| **R Temporo-Occipital** | **247** | | | | |
| posterior MTG | | 4.08 | 45 | -63 | 21 |
| middle OG | | 3.96 | 45 | -75 | 18 |

*The table shows up to three local maxima per cluster more than 8.0 mm apart. L= left; R= right. TP= temporal pole; MTG= middle temporal gyrus; OG= occipital gyrus; ITG= inferior temporal gyrus;*

***Supplementary Table R5*** Significant activation clusters in the mentalising > pain contrast (p < .05, FWE corrected with k= 163 extent threshold and a cluster defining threshold of p< .001, uncorrected)

| **Cluster Name and Location of Maxima** | **Cluster Extent (voxels)** | **Peak (Z)** | **MNI Coordinates (mm)** | | |
|---|---|---|---|---|---|
| | | | **x** | **y** | **z** |
| **L Posterior Medial** | **1990** | | | | |
| precuneus | | 6.2 | -6 | -60 | 21 |
| posterior inferior cingulum | | 5.67 | -9 | -45 | 0 |
| posterior cingulum | | 5.51 | -12 | -45 | 21 |
| **L MTG** | **378** | | | | |
| posterior MTG | | 5.6 | -42 | -63 | 24 |
| posterior MTG | | 3.56 | -54 | -72 | 21 |
| **L Superior Frontal** | **730** | | | | |
| middle SFG | | 4.99 | -18 | 57 | 24 |
| middle SFG | | 4.73 | -27 | 27 | 42 |
| middle MFG | | 4.72 | -33 | 21 | 39 |
| **R MTG** | **163** | | | | |
| AG | | 4 | 42 | -54 | 33 |
| posterior MTG | | 3.49 | 42 | -66 | 21 |

*The table shows up to three local maxima per cluster more than 8.0 mm apart. L= left; R= right; MTG= middle temporal gyrus, SFG= superior frontal gyrus; MFG= middle frontal gyrus; AG= angular gyrus.*

***Supplementary Table R6*** Significant activation clusters representing a three-way conjunction in the interaction judgement > speed judgement Λ false beliefs > photographs Λ mentalizing > pain contrasts (all three contrasts independently treated to a voxel-height threshold of p< .001, uncorrected).

| **Cluster Name and Location of Maxima** | **Cluster Extent (voxels)** | **Peak (Z)** | **MNI Coordinates (mm)** | | |
|---|---|---|---|---|---|
| | | | x | y | z |
| **L inferior ATL** | **92** | | | | |
| lateral anterior ITS / ITG | | 4.37 | -57 | -3 | -33 |
| lateral anterior ITG | | 4.36 | -51 | 3 | -39 |
| **L temporo – parieto – occipital** | **75** | | | | |
| posterior MTG / AG | | 4.14 | -48 | -60 | 15 |
| **L superior frontal lobe** | **75** | | | | |
| middle medial SFG | | 3.92 | -9 | 63 | 27 |
| middle medial SFG | | 3.89 | -9 | 51 | 39 |
| **L precuneus** | **57** | | | | |
| posterior inferior CG | | 3.76 | -3 | -42 | 15 |
| inferior precuneus | | 3.61 | -3 | -51 | 9 |
| inferior precuneus | | 3.31 | -6 | -57 | 18 |
| **R temporo – parieto – occipital** | **50** | | | | |
| posterior MTG / AG | | 3.72 | 45 | -60 | 18 |
| posterior MTG / AG | | 3.65 | 39 | -66 | 21 |

*The table shows up to 3 local maxima per cluster more than 8.0 mm apart. Only clusters greater than 50 voxels reported. L= left; R= right; ITS= inferior temporal sulcus; ITG= inferior temporal gyrus; MTG= middle temporal gyrus; AG= angular gyrus; SFG= superior frontal gyrus; CG= cingulate gyrus.*

***Supplementary Table R7*** Significant activation clusters in the semantic judgement > perceptual judgement contrast (p < .05, FWE corrected with k= 288 extent threshold and a cluster defining threshold of p< .001, uncorrected).

| **Cluster Name and Location of Maxima** | **Cluster Extent (voxels)** | **Peak (Z)** | **MNI Coordinates (mm)** | | |
|---|---|---|---|---|---|
| | | | x | y | z |
| **L temporal – cerebellar** | **2326** | | | | |
| medial posterior MTG | | 5.99 | -42 | -48 | -3 |
| cerebellum | | 5.61 | -42 | -48 | -30 |
| posterior ITS | | 5.2 | -51 | -60 | -6 |
| **R occipital – temporal – cerebellar** | **1933** | | | | |
| inferior occipital gyrus | | 5.92 | 42 | -75 | -6 |
| cerebellum | | 5.36 | 33 | -39 | -27 |
| posterior FG | | 5.32 | 36 | -48 | -24 |
| **L inferior and orbital frontal** | **744** | | | | |
| medial IFG | | 5.7 | -39 | 27 | 15 |
| IFG pars orbitalis | | 5.03 | -45 | 36 | -15 |
| medial IFG pars orbitalis | | 4.42 | -45 | 42 | -3 |
| **L superior frontal** | **288** | | | | |
| middle SFG | | 4.91 | -9 | 54 | 39 |
| posterior SFG | | 3.59 | -12 | 36 | 57 |
| anterior medial SFG | | 3.57 | -3 | 66 | 15 |

*The table shows up to three local maxima per cluster more than 8.0 mm apart. L= left; R= right; MTG= middle temporal gyrus, ITS= inferior temporal sulcus; FG= fusiform gyrus; IFG = inferior frontal gyrus; SFG= superior frontal gyrus;*

***Supplementary Table R8*** Significant activation clusters representing a two-way conjunction in the interaction judgement > speed judgement Λ semantic judgement > perceptual judgements contrasts (the two contrasts independently treated to a voxel-height threshold of p< .001, uncorrected).

| **Cluster Name and Location of Maxima** | **Cluster Extent (voxels)** | **Peak (Z)** | **MNI Coordinates (mm)** | | |
|---|---|---|---|---|---|
| | | | **x** | **y** | **z** |
| **L inferior posterior temporal lobe** | **784** | | | | |
| lateral inferior temporopolar gyrus | | 5.42 | -51 | -48 | -21 |
| posterior FG | | 5.21 | -48 | -57 | -6 |
| cerebellum | | 4.75 | -48 | -66 | -27 |
| **L superior frontal lobe** | **210** | | | | |
| middle SFG | | 5.16 | -12 | 57 | 36 |
| middle SFG | | 4.14 | -3 | 63 | 24 |
| middle SFG | | 3.67 | -18 | 39 | 48 |
| **R inferior posterior temporal lobe** | **812** | | | | |
| inferior occipital gyrus | | 5.01 | 33 | -75 | 0 |
| ITG / FG | | 5.01 | 45 | -60 | -12 |
| FG | | 4.23 | 45 | -54 | -24 |
| **L inferior anterior temporal lobe** | **114** | | | | |
| inferior ATL | | 4.65 | -51 | -6 | -33 |
| left lingual gyrus | **12** | 3.53 | -39 | -84 | -18 |
| **L temporal pole** | **32** | | | | |
| superior TP | | 3.5 | -42 | 24 | -33 |
| superior TP | | 3.48 | -27 | 18 | -33 |
| left gyrus rectus | **24** | 3.47 | 0 | 42 | -24 |

*The table shows up to 3 local maxima per cluster more than 8.0 mm apart. Only clusters greater than 50 voxels reported. L= left; R= right; FG= fusiform gyrus; SFG= superior frontal gyrus; ITG= inferior temporal gyrus; ATL= anterior temporal lobe; TP = temporal pole*
